# Regionalisation of muscle nucleus heterogeneity in foetal skeletal muscle

**DOI:** 10.64898/2026.10.29.762763

**Authors:** Tamaki Kurosawa, Cédrine Blavet, Marie-Ange Bonnin, Tarek Gharsalli, Marie-Claire Delfini, Estelle Hirsinger, Delphine Duprez

## Abstract

Skeletal muscle is organized into functional domains with attachments at both extremities to tendons via the myotendinous junction (MTJ) and innervation in the middle via the neuromuscular junction (NMJ). In adult muscle, the molecular heterogeneity of muscle domains is described at the level of myonuclei with specialized MTJ and NMJ myonuclei at the tips and middle of muscle fibre. However, the regionalisation of molecular heterogeneity remains poorly understood during development. Combining single-cell and single-nucleus RNA sequencing approaches with *in situ* hybridization experiments to limb muscles of chicken embryos, we uncover transcriptionally distinct muscle nuclei according to tip and central domains, in both myofibres and mononucleated myogenic cells. At early foetal stage, all myonuclei display either a tip or a central domain identity. Immature mononucleated myogenic cells are enriched in the tip domain. Tip myonuclei of foetal muscles show a robust transcriptional “signal transduction” signature, which highlights molecular crosstalks with adjacent tendon fibroblasts but also with adjacent immature myogenic cells. We propose that the tip muscle domain is a signalling centre interacting with tendon fibroblasts to regulate muscle/tendon attachment and with adjacent mononucleated muscle cells to regulate myogenesis in limb foetal muscles.

**Summary statement:** Myonuclei and mononucleated muscle cells exhibit molecular regionalisation according to tip and central domains in foetal skeletal muscle

## Introduction

Skeletal muscle is composed of multinucleated muscle fibres surrounded by mononucleated myogenic cells in addition to numerous other cell types including, fibroblasts, blood vessels, immune cells and neuronal projections. Muscle fibres compose the functional unit of muscle allowing muscle contraction involved in locomotion, posture and feeding. In adult muscle, multinucleated muscle fibres are composed up to 800 (mouse) to 40 000 (human) nuclei named myonuclei (Cramer et al., 2020; Hansson et al., 2020). Each limb muscle is unique with specific characteristics, such as location, shape, size and fibre types. Despite this diversity, all muscles display the same type of organization with tendon attachments at both extremities via myotendinous junction (MTJ) and innervation targeting the middle of the muscle via neuromuscular junction (NMJ). This defines the tip and central functional muscle domains. The molecular heterogeneity of the functional domains has been established at the level of the muscle fibre in 2020 with single-nucleus RNAsequencing strategy to capture transcriptomic profiles of myonuclei (Dos Santos et al., 2020; Kim et al., 2020; Petrany et al., 2020). In adult muscles, myonuclei display specific transcriptional profiles associated to MTJ (tip) and NMJ (central), reviewed in (Korb et al., 2024). In adult muscle, the specialized MTJ and NMJ myonuclei account for less than 10% of all myonuclei (in mouse limb skeletal muscles), the majority being the body myonuclei displaying no specific identity other than a molecular signature of contractile muscle proteins, reviewed in (Korb et al., 2024). Molecular markers have been identified for the specialized MTJ and NMJ myonuclei of adult muscles. *COL22A1* is a recognized marker of MTJ myonuclei located at the tips of muscle fibres close to tendons, in human and mouse adult skeletal muscles, reviewed in (Mackey, 2025). The postsynaptic components of NMJ are recognized to be transcribed in subsynaptic myonuclei adjacent to motoneurons targeting the middle of adult skeletal muscle (Ham et al., 2025; Kim et al., 2020).

While MTJ and NMJ are mature after birth, MTJ and NMJ markers are already expressed in muscle during development. The key MTJ marker, *Col22a1*/Collagen XXII is regionalised (at both transcript and protein levels) in tip myonuclei of developing muscles of mouse, chicken and zebrafish embryos (Charvet et al., 2013; Koch et al., 2004; Malbouyres et al., 2022; Nicolas et al., 2024). There are also sporadic data with description of genes specifically located at the tip of foetal muscles, such as *FGF4, FGF6, ANKRD1, CTGF, ID2* in chicken and mouse embryos (Armand et al., 2006; deLapeyrière et al., 1993; Esteves de Lima et al., 2021; Esteves de Lima et al., 2016). Key post-synaptic NMJ genes are expressed during development, however with large central expression domains according to the pre-patterning paradigm (Kummer et al., 2006; Li et al., 2018; Tintignac et al., 2015).

Mononucleated muscle cells have been recognized to be molecularly heterogeneous according to embryonic and foetal myogenesis (Hutcheson et al., 2009; Murphy and Kardon, 2011) (Messina et al., 2010; Toulouse et al., 2025). The embryonic and foetal PAX7+ progenitors and *MYOD*+/MYOD+ myoblasts appeared to be homogenously distributed in chicken and mouse limb foetal muscles with no obvious regionalisation according to tip and central muscle domains (Esteves de Lima et al., 2022; Esteves de Lima et al., 2016; Murphy and Kardon, 2011; Toulouse et al., 2025). In contrast, the fusion-associated genes, myomaker and myomixer, and myomixer protein are concentrated in the middle of foetal muscles in both myonuclei and mononucleated muscle cells pointing to fusion events in the middle of muscle and excluded from the tip domain during development (Esteves de Lima et al., 2022; Nahlé et al., 2025; Nicolas et al., 2024).

Although the molecular heterogeneity is well established at the level of adult muscle fibres according to central and tip functional domains, the transcriptional heterogeneity of developmental myonuclei is less understood. The molecular heterogeneity of mononucleated myoblasts within developing muscles according to central and tip domains is still an open question. How this molecular heterogeneity coordinates with myogenic progression is also not understood. Using RNA sequencing approaches at the level of the nucleus/cell combined with extensive in situ hybridization experiments to limb foetal muscles, we mapped the molecular heterogeneity of muscle nuclei at the level of myonuclei and mononucleated cells according to muscle domains.

## Results

### Single-nucleus RNA-sequencing of limb foetal muscles illustrates myogenic progression and reveals two populations of myonuclei

In order to assess the molecular landscape of foetal myonuclei according to central and tip domains, we turned to single-nucleus RNA-sequencing (snRNAseq) strategy of chicken limb muscles. We generated snRNAseq dataset on whole chicken forelimbs at E10 foetal stage (**Fig. S1).** Clustering analysis showed that skeletal muscle nuclei segregate from other limb cell types into three clusters, cluster 6,7,10 (**Fig. S1)**. To analyse skeletal muscle nuclei, we bio-informatically extracted nuclei of clusters 6,7,10 from the whole limb dataset. Unsupervised clustering analysis of the 507 muscle nuclei at resolution 0.5 revealed five distinct clusters (**Figs. 1, S2**) associated with unfiltered (**Table S1**) and filtered (pct.1>0.40 and pct.2<0.25, **Table S2**) marker lists. Each cluster can be identified with the expression of classical/recognized muscle markers (**Fig. 1A-D**). Normalized expression of muscle markers visualised with violin plots (**Fig. 1C)** and feature plots **(Fig. S2B**) allowed us to identify cluster 0 as *PAX3*+ *FGFR4*+ muscle progenitors. The expression of *FGFR4, MYOD, TMEM8C* was enriched in cluster 1, identifying myoblasts entering into the myogenic program. Nuclei of cluster 3 were enriched with the expression of *MYOG, TMEM8C*, *MYH1B* and *MYH1G* reflecting one step further into muscle differentiation and pointing to myocytes **(Figs. 1C, S2B**). Clusters 2 and 4 displayed a signature of myonuclei (the nuclei of multinucleated muscle fibres) with the complete absence of *FGFR4* expression and the expression of *MYH7B, TNNT1* and *TNNT3* as top differentiation genes of the filtered marker list for clusters 2 and 4 (**Figs. 1C, S2B,C, Table S2).** In particular, *MYH7B* was exclusive to the two myonuclei clusters (**Figs. 1C, S2B**).

**Figure 1.**
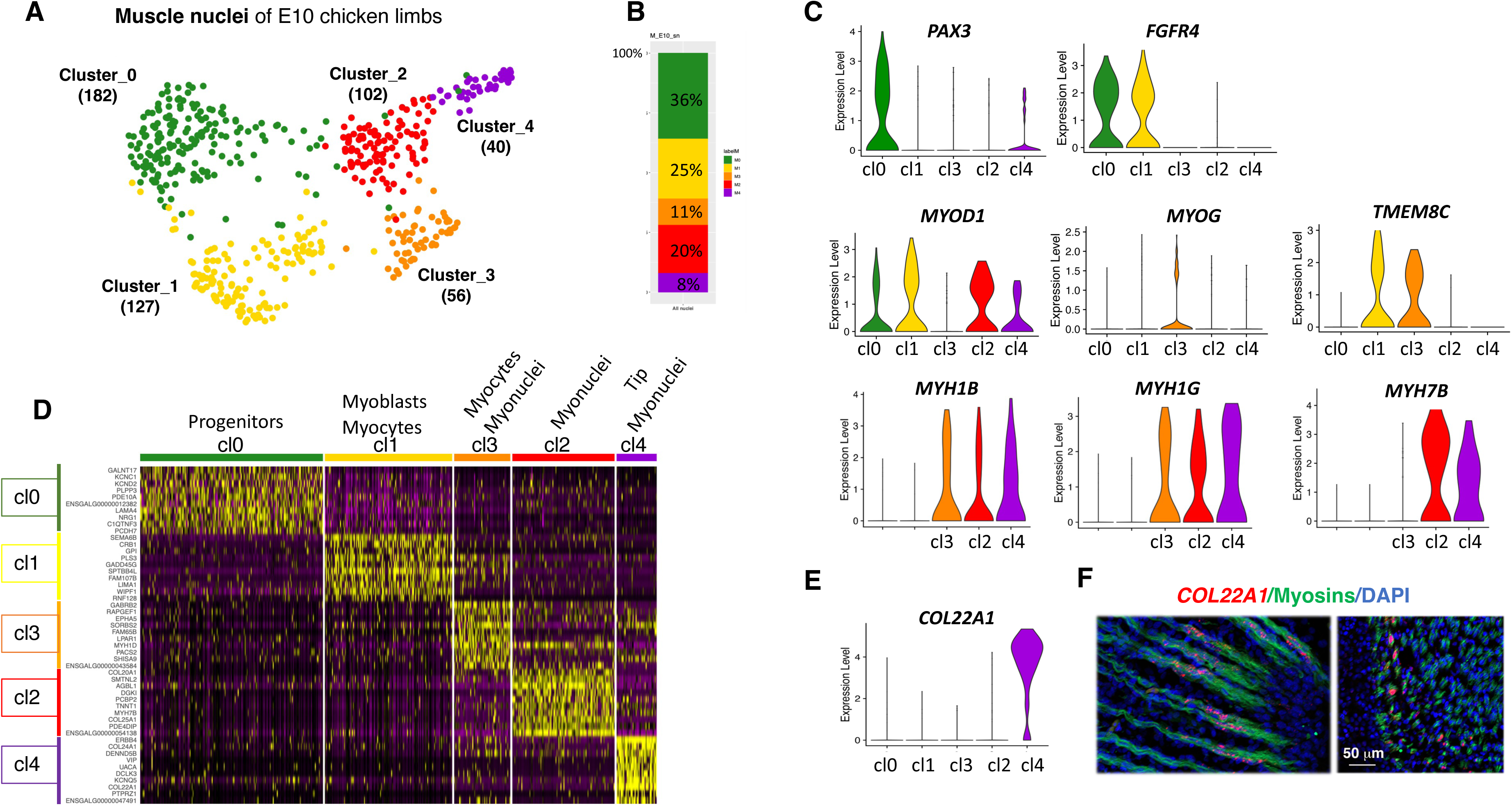
Clustering analysis of the E10 limb muscle nuclei illustrates myogenic progression and identifies two clusters of myonuclei. **(A)** UMAP plot showing the distribution of the five muscle clusters and their cluster size, cluster 0 (182 nuclei), cluster 1 (127 nuclei), cluster 2 (102 nuclei), cluster 3 (56 nuclei) and cluster 4 (40 nuclei), in the muscle snRNAseq dataset from E10 chicken limbs. **(B)** Stacked histogram showing the percentage of nuclei per cluster in the E10 muscle snRNAseq datasets. **(C)** Violin plots showing the log-normalized expression levels for the following muscle markers (ordered according to myogenic progression), *PAX3*, *FGFR4, MYOD1, MYOG, TMEM8C*, *MYH1B*, *MYH1G, MYH7B* across clusters. **(D)** Heatmap showing the relative expression of the top 10 markers across all nuclei of the five clusters, ordered according to myogenic progression. Upregulated genes are in yellow, while downregulated genes are in purple. **(E)** Violin plot showing the log-normalized expression levels for *COL22A1* across clusters. **(F)** Fluorescent in situ hybridization to longitudinal and transverse muscle sections from E10 chicken limbs with *COL22A1* probe (red) combined with myosin immunolabelling (green) and DAPI staining (blue). Images focused on tip muscle regions.

The overlapping expression of the key muscle markers (*PAX3, FGFR4, MYOD, TMEM8C, MYOG, MYHs, TNNTs*) across clusters (**Figs. 1C, S2B,C**) was consistent with the overlapping expression of these markers along the myogenic progression, reviewed in (Comai and Tajbakhsh, 2014; Esteves de Lima and Relaix, 2021). The heatmap and dotplots of the top 10 markers of each cluster (ordered according to myogenic progression) showed a specific molecular signature for each cluster despite overlapping expression of muscle markers across clusters (**Figs. 1D, S2A**). As expected, the snRNAseq provide similar information as the scRNAseq (Esteves de Lima et al., 2022) concerning the myogenic progression from *PAX7*+/*FGFR4*+ progenitors (clusters 0 for both datasets), the differentiating *MYOD1* state (cluster 1 for both datasets) to the myosin/troponin differentiation state (cluster 3 for snRNAseq and cluster 2 for scRNAseq) (**Figs. 1, S2, S3**). The proportion of the three populations (muscle progenitors, MYOD1 state and myosin/troponin state) was comparable between snRNAseq and scRNAseq datasets (**Fig. S3B,C**). This indicates that both snRNAseq and scRNAseq were valid strategies to visualize myogenic progression up to the differentiation step of mononucleated cells. However, only the snRNAseq strategy allowed us to capture the myonucleus landscape of foetal muscles and identify two populations of myonuclei in limb foetal muscles of E10 chicken embryos.

We conclude that the snRNAseq dataset of limb foetal muscle illustrates myogenic progression from muscle progenitors to multinucleated cells, and identifies two populations of myonuclei.

### Foetal myonuclei located at muscle tips exhibit a robust transcriptional signature sharing molecular features with adult MTJ

In addition to myosin and troponin gene expression, myonuclei of cluster 4 exhibited a clear identity characterised by the exclusive expression of the MTJ marker *COL22A1* in 90% of cluster 4 myonuclei (**Figs. 1E, S4, Table S2**). *COL22A1* expression was restricted to tip myonuclei of limb foetal muscles (**Fig. 1F**); this regionalised expression being concomitant with myotube formation during chicken development (**Figs. S4 E-J**). To further characterize tip myonucleus identity, we analysed the cluster 4-associated markers. The long marker lists of cluster 4 (464 and 118 genes for the non-filtered and filtered list, respectively) indicated a strong molecular identity for cluster 4 myonuclei (**Tables S1,S2)**. Numerous genes with high fold-changes were unique marker of cluster 4 and displayed high pct.1 and low pct.2 (**Tables S1,S2)**. Moreover, the cluster 4 was detected at resolution 0.4 and did not change up to resolution 0.9 (**Fig. S4A,B**). To select for the most specific markers, we chose the top six genes of cluster 4 filtered marker list (ordered by fold-change) fulfilling the following criteria: pct.1 greater than 0.8, pct.2 lower than 0.2 and gene exclusively associated to cluster 4 (**Table S2**) and performed FISH experiments to the selected genes. *COL22A1* was the top of these six genes (log2FC_5.2, pct1_0.9, pct2_0.07), followed by *ERBB4* (Erb-B2 receptor tyrosine kinase 4) coding for the tyrosine kinase receptor ERBB4 (pct1_1, pct2_0.12), *DCLK3* (DoubleCortin-Like Kinase 3) coding for a serine/threonine kinase (pct1_0.8, pct2_0.04), *COL24A1* coding for the alpha1 chain of type XXIV collagen (pct1_0.9, pct2_0.17), *VIP* (Vasoactive Intestinal Peptide) coding for a peptide hormone (pct1_0.8, pct2_0.03) and *UACA* gene coding for a neuropeptide *UACA* (Uveal Autoantigen with Coiled-coil domains and Ankyrin repeats) (pct1_0.8, pct2_0.06). These genes showed high fold-changes, with log2FC ranging from 5.2 to 3.5. Remarkably, the six selected genes exhibited a selective expression in tip myonuclei of myotubes close to the muscle/tendon interface of limb foetal muscles (**Figs. 2A-J, S5**). In addition to *DCKL3*, we identified another member of doublecortin family (down the cluster 4 marker list with avg_log2FC 2,4, pct1 0.75, pct2 0.07), *DCX* (DoubleCortin) as being expressed in tip myonuclei close to tendon visualised with *COL12A1* (**Fig. S5C-D**). We also validated two other genes (further down the cluster 4 marker lists) with *SMAD9* and *CXCR7* as being also expressed in tip myonuclei (**Fig. 2H-N, Tables S1,S2**). The selective expression of *SMAD9* at the tips of foetal muscles was consistent with the location of pSMAD1/5/9 in tip myonuclei of chicken foetal limb muscles (Esteves de Lima et al., 2021). Tip myonuclei appeared to be molecularly homogeneous. 57.5% of tip myonuclei expressed the combination of the top six markers (*COL22A1, ERBB4, DCLK3, COL24A1, UACA, VIP*) and 100% of them expressed a combination of at least 2 markers (**Fig. S5H**). Genes already described as being expressed in tip myonuclei of limb foetal muscles were present in the marker lists of cluster 4 (**Tables S1,S2**), such as *FGF4* (Edom-Vovard et al., 2002), *FGF6* (deLapeyrière et al., 1993), *MEF2C* (Nicolas et al., 2024), *ID2* (Esteves de Lima et al., 2021), and *ANKRD1* (Esteves de Lima et al., 2016). Cluster 4 therefore reveals the molecular signature of myonuclei located at the tip of foetal myofibers.

**Figure 2.**
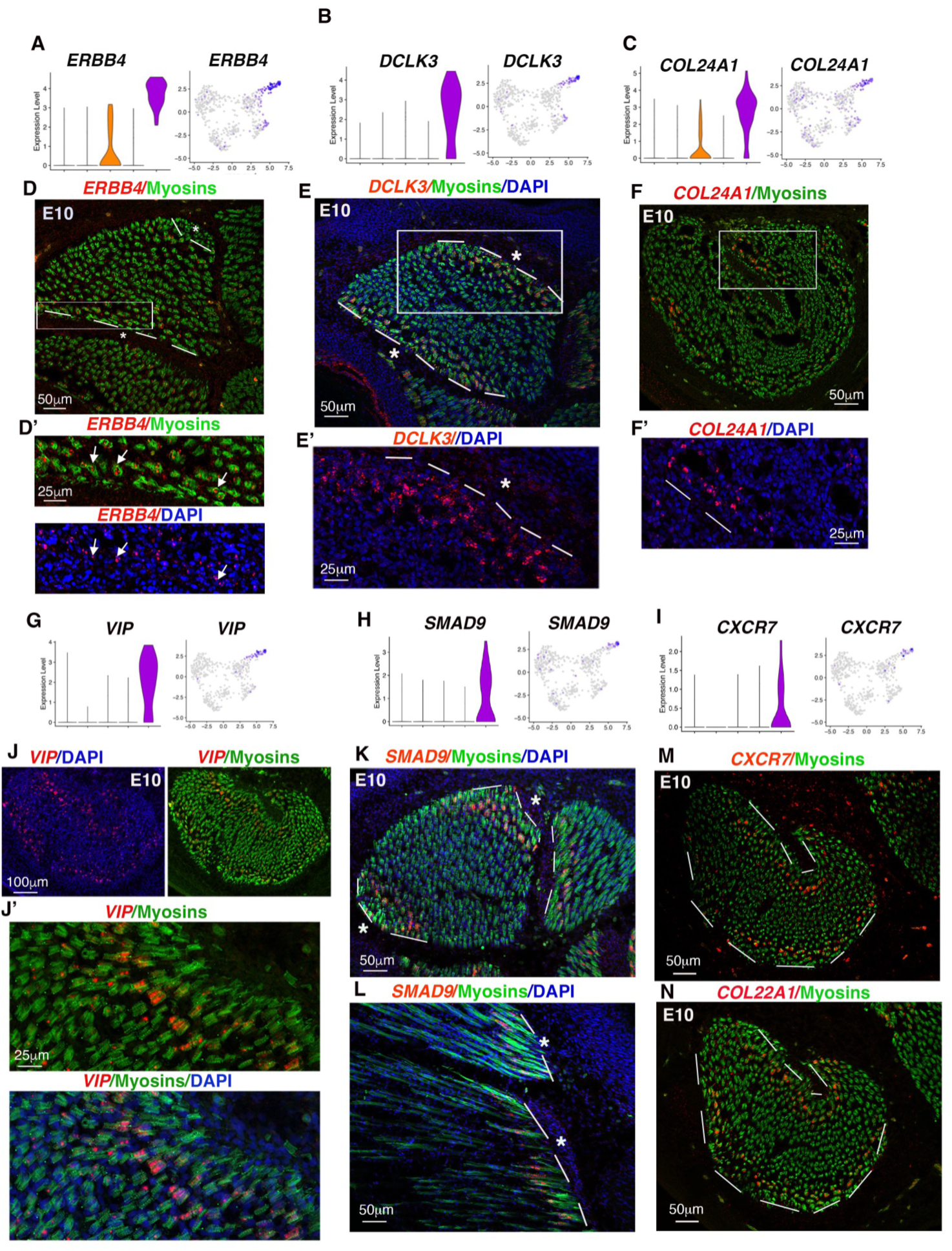
Molecular markers for tip myonuclei of limb foetal muscles. (**A-C**) Violin and feature plots showing the log-normalized expression levels and the distribution of marker+nuclei across muscle clusters and nuclei in the E10 muscle snRNAseq dataset for *ERBB4* (A), *DCLK3* (B), *COL24A1* (C) genes. (**D-F**) Fluorescent in situ hybridization to limb muscle transverse sections of chicken embryos at E10 with *ERBB4* (D,D’), *DCLK3* (E,E’), *COL24A1* (F,F’) probes (red), followed with immunohistochemistry with the MF20 antibody recognising myosins (green) and DAPI staining (blue). **(G-I)** Violin and feature plots showing the log-normalized expression levels and the distribution of marker+nuclei across muscle clusters and nuclei in the E10 muscle snRNAseq dataset for *VIP* (G), *SMAD9* (H) and *CXCR7* (I) genes. (**J-N**) Fluorescent in situ hybridization to transverse (J,K,M,N) and longitudinal (L) limb muscle sections of E10 chicken embryos with the *VIP* (J,J’), SMAD9 (K,L), CXCR7 (M) and COL22A1 (N) probes (red), followed with an immunohistochemistry with the MF20 antibody recognising myosins (green) and DAPI staining (blue). All these cluster 4 marker genes, *ERBB4, DCLK3, COL24A1, VIP, SMAD9* and *CXCR7* displayed a regionalised expression in tip myonuclei of limb foetal muscles. Dashed lines indicate the muscle/tendon interface. Stars indicate tendon location.

The molecular signature of foetal tip myonuclei exhibited similarities with adult MTJ myonuclei (**Fig. S6)**. Among the six common MTJ myonuclei markers identified in five studies of adult human and mouse muscles (Karlsen et al., 2023), two genes (*COL22A1*, *SORBS2*) were present in the marker lists of cluster 4 of chicken foetal muscles (**Tables S1, S2**). In addition, by crossing the marker lists of adult MTJ clusters from human muscle snRNAseq datasets (Karlsen et al., 2023) with the cluster 4 unfiltered list of foetal tip myonuclei from chicken muscle snRNAseq dataset, we found that 47 genes (7%) were shared genes (**Fig. S6A-C**). Five shared genes (*COL22A1*, *SMAD9, FGF6, ANKRD1* and *MYH15*) (**Fig. S6D**) were validated as being expressed in tip myonuclei by FISH in developing chicken limb muscles (**Figs 1F**, **2K,L**, **3F**).

**Figure 3.**
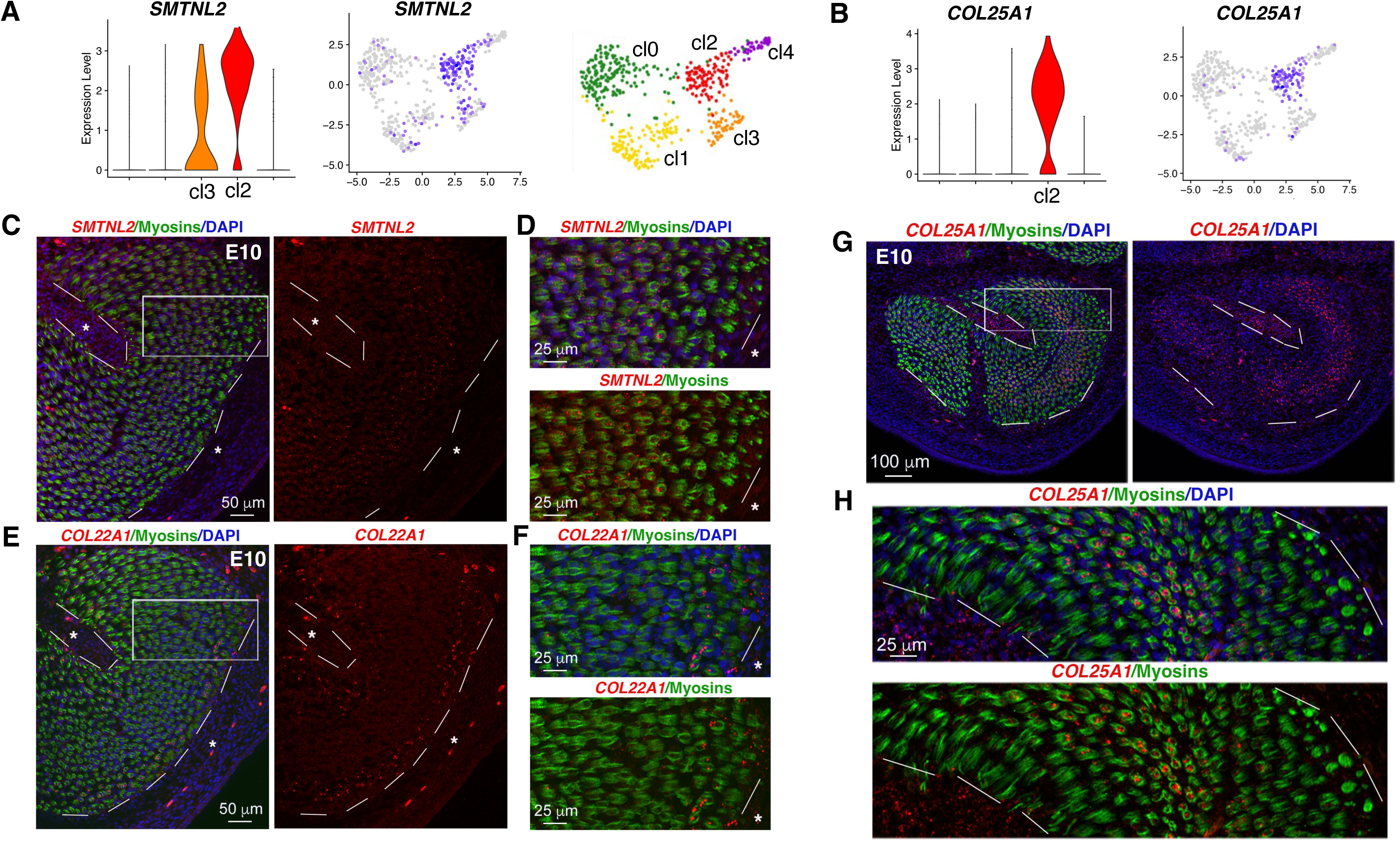
SMTNL2 and *COL25A1* are expressed in central myonuclei of limb foetal muscles. (**A,B**) Violin and feature plots showing the log-normalized expression levels and the distribution of *SMTNL2* (A) and *COL25A1* (B), cluster 2 markers, across clusters and nuclei, in the E10 muscle snRNAseq dataset. Repetition of the UMAP plot coloured by clusters, shown in Fig. 1A. (**C-F**) Fluorescent in situ hybridization to adjacent and transverse muscle sections with *SMTNL2* (C,D) and *COL22A1* (E,F) probes (red) followed with an immunostaining for myosins using the MF20 antibody (green) and DAPI staining. (D,F) are high magnifications of the squared muscle areas in (C,E), respectively. *SMTNL2* is expressed in central myonuclei and not in tip myonuclei labelled with *COL22A1*. (**G,H**) Fluorescent in situ hybridization to limb muscle transverse sections with *COL25A1* probe (red) followed with an immunostaining for myosins using the MF20 antibody (green) and DAPI staining. (H) is a high magnification of the squared muscle area in (G) showing *COL25A1*/Myosins/DAPI and *COL25A1*/Myosins staining. In addition to being expressed in central myonuclei, *COL25A1* is also weakly expressed in tendons. Dashed lines show muscle/tendon interfaces. White stars indicate tendon locations.

We conclude that myonuclei located in the tip domain of chicken limb muscles exhibit a robust molecular identity at foetal stage and share molecular features with adult MTJ myonuclei.

### Centrally-located myonuclei display a NMJ signature

To determine the molecular identity of cluster 2 myonuclei, we selected markers exclusively associated with cluster 2 from the filtered marker list (**Table S2**). We were able to identify *STMNL2* and *COL25A1* as specific cluster 2 markers, detected in 90% and 82% of cluster 2 myonuclei, respectively (**Fig. 3A,B, Table S2)**. *STMNL2* and *COL25A1* genes were specifically expressed in central myonuclei of limb foetal muscles and excluded from tip myonuclei expressing *COL22A1* (**Fig. 3C-H**). Despite being only listed in the cluster 2 marker list, *STMNL2* was also slightly observed in cluster 3 in violin and feature plots (**Fig. 3A**). *STMNL2* code for a smoothelin-like protein not well characterized, but *STMNL1* is associated with the regulation of muscle cell contractility following exercise training (Murali and MacDonald, 2018). *COL25A1* gene codes for type COLXXV, a transmembrane collagen involved in NMJ formation during mouse limb development (Munezane et al., 2019; Tanaka et al., 2014). Key post-synaptic NMJ genes, such as *VAV3, LRP4, CHRNG, CAMK4* (ordered by fold-changes), but not *CHRNA1* and *ETV5,* were listed in cluster 2 unfiltered marker list (**Table S1**). Nonetheless, all the above genes were detected in cluster 2, albeit not exclusively as they were also detected in nuclei of mononucleated cells (**Fig. S7A,B).** In the case of *CHRNA1* and *ETV5,* this was consistent with their *in situ* location where they were expressed in the central domain of muscle, in a pattern complementary to *FGF4* expression at muscle tips (**Fig. S7C**). Despite the fact that the master gene for fusion also displayed a central location in developing muscle (Esteves de Lima et al., 2022), *TMEM8C* was not listed in cluster 2 marker lists (**Tables S1, S2**), indicating that *TMEM8C* transcripts were mainly located in mononucleated muscle cells in the central domain.

We conclude that cluster 2 identifies central myonuclei that harbour a NMJ molecular signature in E10 foetal muscles.

### Tip and central myonucleus markers cover all myonuclei of early foetal muscles

The expression domains of tip (*COL22A1*) and central (*COL25A1* and *STMNL2*) myonuclei markers appeared to be complementary at early foetal stage (**Fig. 3**). Moreover, the expression domains of the tip myonuclei marker (*FGF4*) appeared to be mutually exclusive with that of the fusion gene *TMEM8C* (Esteves de Lima et al., 2022). The central expression domain of the *CHRNA1* NMJ marker was similar to that of the *TMEM8C* fusion gene and mutually exclusive from that of *COL22A1* (**Fig. 4A,B**). In order to have a cellular resolution we performed HCR FISH with tip and central myonucleus markers (**Fig. 4C-F**). Surprisingly, we observed no gap between tip myonuclei labelled with *COL22A1* and central myonuclei labelled with *COL25A1* in transverse and longitudinal sections of foetal muscles (**Fig. 4C,D**). We observed very little gap between the tip myonuclei labelled with *COL22A1* and the central expression domains of *CHRNA1* (NMJ) and *TMEM8C* (fusion) genes, domains that include myonuclei and mononucleated cells (**Fig. 4E-F).** The absence of non-labelled myonuclei in the double labelling tip (*COL22A1*) and central (*COL25A1*) markers indicated the absence of non-specialized myonuclei (body myonuclei) in E10 foetal muscles. The body myonuclei were detected in E14 foetal muscles with the narrowing of the specialized myonuclei domains, assessed with the MTJ (*COL22A1*) and NMJ (*CHRNA1*) gene expression (**Fig. 4G,H**). However, the central *TMEM8C* and *COL25A1* expression domains showed no obvious narrowing (**Fig. 4I,J**), indicating that fusion events occurred in central region independently of NMJ maturation.

**Figure 4.**
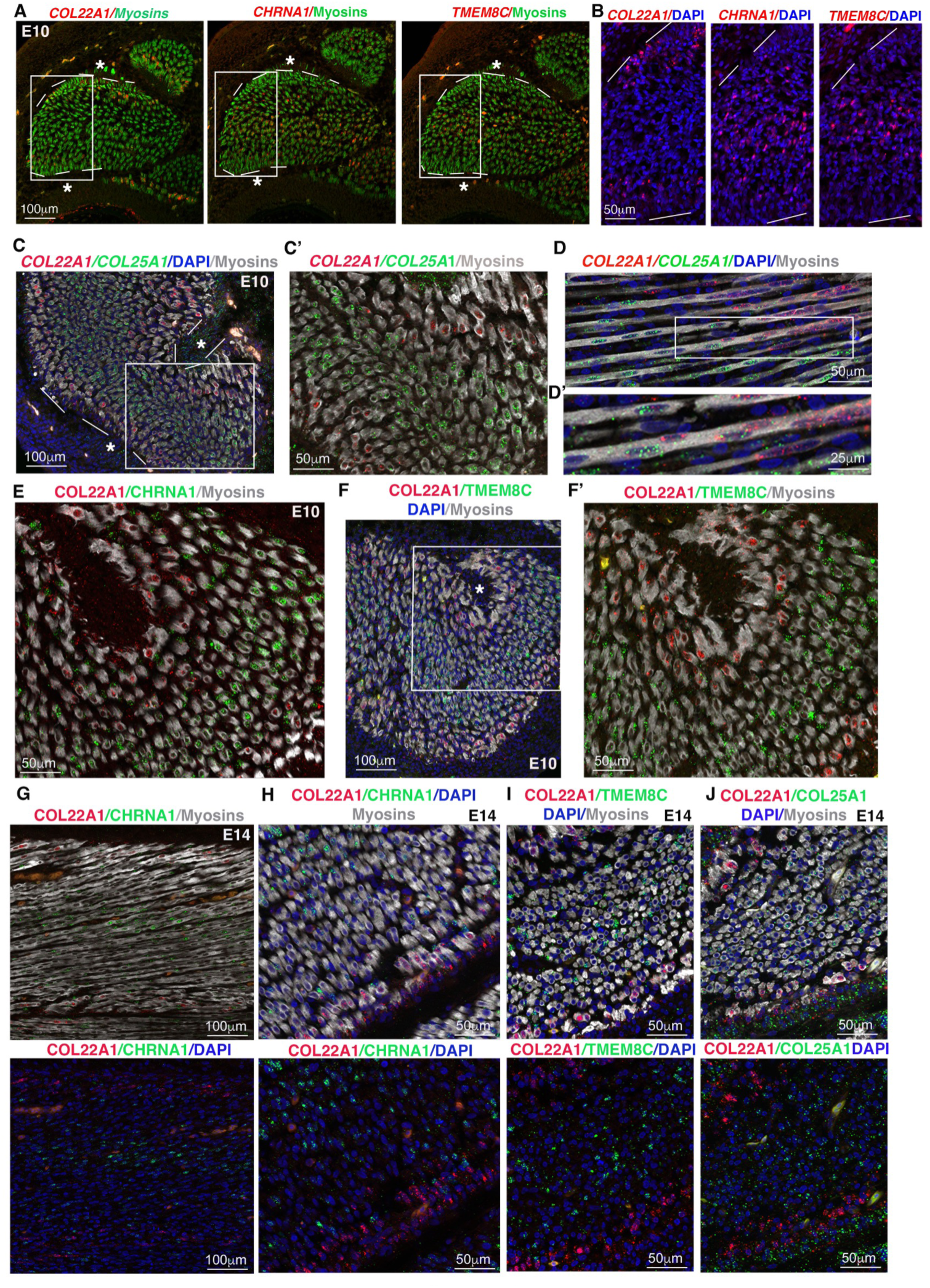
Myonuclei express either tip or central markers at early foetal stage. (**A,B**) Fluorescent in situ hybridization to adjacent limb muscle transverse sections at E10 with COL22A1, CHRNA1 and TMEM8C probes (red) followed with an immunostaining for myosins using the MF20 antibody (green) and DAPI staining. (A) Low magnifications of a dorsal muscle showing the tip expression domain of *COL22A1* and the central expression domains of the postsynaptic gene *CHRNA1* and fusion gene *TMEM8C.* (B) are high magnifications of the squared regions in (A) showing the regionalisation of *COL22A1, TMEM8C* and *CHRNA1* genes. (**C-D’**) Double HCR-FISH to transverse (C,C’) and longitudinal (D,D’) limb muscle sections at E10 with the following COL22A1(red) / COL25A1 (green) probes followed with an immunostaining to label myosins (grey). (**E**) Double HCR-FISH to transverse limb muscle sections at E10 with the following COL22A1(red) / CHRNA1(green) probes followed with an immunostaining to label myosins (grey). (**F,F’**) Double HCR-FISH to transverse limb muscle sections at E10 with the following COL22A1(red) / TMEM8C (green) probes followed with an immunostaining to label myosins (grey). (**G-J**) Double HCR-FISH to longitudinal (G) and transverse (H-J) limb muscle sections at E14 with the following COL22A1(red) / CHRNA1(green) (G,H), COL22A1(red) / TMEM8C (green) (I), COL22A1(red) / COL25A1 (green) (J) probes followed with an immunostaining to label myosins (grey). (G-J) Top and bottom panels show the same section with different labelling. Dashed lines show muscle/tendon interfaces. White stars indicate tendon locations.

We conclude that specialized MTJ and NMJ myonuclei cover tip and central domains in limb muscles at early E10 foetal stage, and display a narrowing at E14 foetal stage.

### Functional identity of foetal tip and central myonuclei

In order to define a functional identity of tip and central myonuclei, we analysed the filtered marker lists of clusters 4 and 2 (**Table S2**) using the Reactome Pathway Database (Fabregat et al., 2017; Wu and Haw, 2017) (**Fig. 5A,B, Table S6**).

**Figure 5.**
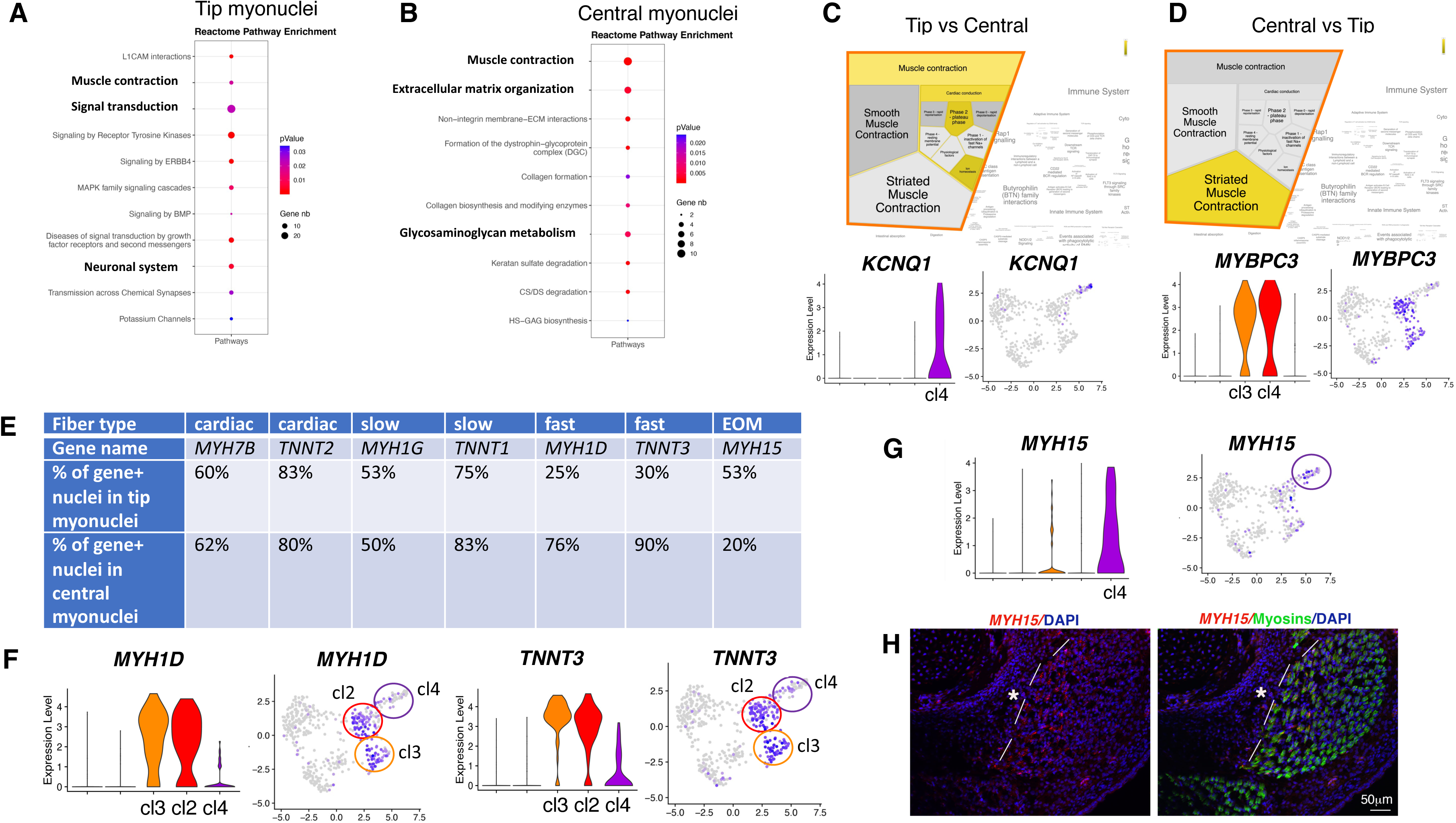
Tip myonuclei display a “signal transduction” identity. (**A,B**) Dotplots showing the p-value and gene number for selected Pathway names issued from the Reactome analysis performed on cluster 4 filtered marker list (A), and on cluster 2 filtered marker list (B). A selection of Pathway names in the Reactome hierarchy with high #Entities found were plotted. (**C,D**) Pathway names (highlighted in yellow) significantly enriched from the Reactome analysis performed on cluster 4 (tip) versus cluster 2 (central) (C), and on cluster 2 (central) versus cluster 4 (tip) (D). Violin and feature plots showing the log-normalized expression levels and the distribution of *KCNQ1* expressed in tip myonuclei (C) and *MYBPC3* enriched in central myonuclei (D). (**E**) Percentage of gene+ nuclei in tip myonuclei (cluster 4) and central myonuclei (cluster 2) for cardiac genes (*MYH7B, TNNT2*), slow muscle genes (*MYH1G, TNNT1*), fast muscle genes (*MYH1D, TNNT3*) and the EOM muscle gene, *MYH15*. **(F)** Violin and feature plots showing the log-normalized expression levels and the distribution of the fast type genes, *MYH1D* and *TNNT3* across clusters and nuclei. **(G)** Violin and feature plots showing the log-normalized expression levels and the distribution of *MYH15+* nuclei across clusters and nuclei. (**H)** Fluorescent in situ hybridization to transverse muscle sections of E9 limbs with *MYH15* probe (red), followed by an immunostaining with the MF20 antibody to label myosins (green) and DAPI staining (blue). Dashed white lines show the muscle/tendon interface and white stars indicate tendon location.

For tip myonuclei, the reactome analysis highlighted a significant enrichment for the following Reactome pathways “L1CAM interactions”, “muscle contraction”, “signal transduction” and “neuronal system” (**Fig. 5A, Table S6**). The “L1CAM interactions” category, that includes *LAMC1*, *NRCAM* and *DCX*, pointed to migration and cell-cell contact mediated by the L1CAM family of cell adhesion molecules. As an example, the microtubule-associated proteins, DCX/DCLK3 (**Figs. 2E, S5F**) are involved in cell-cell adhesion mediated migration during neurogenesis (Ayanlaja et al., 2017). The neuronal system with “Potassium channels” categories pointed to the excitation-contraction coupling regulation, as exampled with the potassium-voltage channels *KCNQ1* and *KCNQ5*. The “signal transduction” category represented the category with the highest number of genes and several sub-categories such as “signaling by Receptor Tyrosine Kinases” (*GALNT3, COL24A1, EGF, VEGFC, GAB2, LAMC1, PTPRZ1, FGF9, ERBB4, RPS6KA2, ID2, HBEGF, MXD4*), “signaling by ERBB4” (*ERBB4, EGF, MXD4, HBEG*), “MAPK family signaling cascades” (*DUSP10, FGF9, ERBB4, EGF, HBEGF*) and “signaling by BMP” (*SMAD9, SMAD7*). This showed that the EGF/ERBB4, FGF/MAPkinase, and BMP/SMAD9 signalling pathways were enriched in tip myonuclei. The ERBB4 receptor is involved in a plethora of biological processes, including a role in cardiac and skeletal muscle differentiation (Fernández-Simón et al., 2026; Jiang et al., 2026), but also a role in neuronal migration during development and adulthood (Fornasari et al., 2016). In addition to EGF, FGF or BMP signalling pathways, the *VIP* gene coding for vasoactive intestinal peptide, was also highly specific to tip myonuclei compared to muscle nuclei (**Fig. 2G,J, Table S2**). VIP is a neuropeptide distributed in central and peripheral nervous systems and involved in neurogenesis and peripheral nerve regeneration (Cochaud et al., 2015; Zaben and Gray, 2013).

The reactome pathways enriched in central myonuclei were completely different from those enriched in tip myonuclei (**Fig. 5A,B**). For central myonuclei, the reactome analysis did not show any “signal transduction” category, but rather highlighted a significant enrichment for, “Extracellular matrix organisation” and “Glycosaminoglycan metabolism » (**Fig. 5B, Table S6**). Genes under the categories of “Extracellular matrix organisation” include genes coding one secreted collagen *(COL20A),* two transmembrane collagens (*COL13A1*, *COL25A1*), and membrane components linking the extracellular matrix to the intracellular cytoskeleton, such as delta-sarcoglycan (*SGCD*), syndecan-2 (*SDC2*) and Syntrophin-γ2 (*SNTG2*). Consistent with their central location in fœtal muscle, these genes were all associated to NMJ formation, structure, maintenance or dysfunction (Aittaleb et al., 2017; Gloriani et al., 2025; Härönen et al., 2019; Hastings et al., 2025; Wakabayashi, 2020; Zhou et al., 2021).

In order to specifically address the difference of molecular signatures between tip and central myonuclei, we analysed the markers enriched in cluster 4 versus cluster 2, and in cluster 2 versus cluster 4 (**Table S7**). In addition to the genes enriched in clusters 4 and 2 versus all the other clusters (**Tables S1,S2**), this analysis highlighted additional genes (**Table S7**). The analysis of these marker lists using the Reactome Pathway Database (**Table S7**, **Fig. 5C,D**) pointed to a “cardiac conduction” feature of muscle contraction for tip myonuclei, with cardiac-specific ion channels/pumps (*KCNQ1, ITPR1, ATP2B2, CACNA1*). Mutations in *KCNQ1* or *CACNA1* genes lead to severe cardiac arrhythmias (Dehghani-Samani et al., 2019). In contrast, central myonuclei displayed a “striated muscle contraction” feature with genes coding for sarcomeric proteins of skeletal muscle (*MYBPC3, TNNT3*), (**Fig. 5C,D).** Because of the different muscle contraction features (**Fig. 5C,D**), we looked for selective genetic myosin program in tip myonuclei versus central myonuclei (**Fig. 5E-H**). While the cardiac myosin genes (*MYH7B, TNNT2*) and the slow myosin genes (*MYH1G, TNNT1*) were expressed similarly in both tip and central myonuclei (**Fig. 5E, Table S2**), tip myonuclei appeared to be depleted in genes involved in the fast fibre type program (*MYH1D, TNNT3*) (**Fig. 5F**). This is indirectly consistent with the gene enrichment of the slow fibre type at the tips of muscle fibres in postnatal mouse muscles (Dos Santos et al., 2022). In addition, the unconventional myosin *MYH15,* expressed in ventricles of chicken adult hearts (Garriock et al., 2005) was enriched in tip myonuclei (**Fig. 5G)** and selectively expressed in tip myonuclei of chicken limb muscles (**Fig. 5H**). *MYH15* is also known to be expressed in rat extraoccular muscles and muscle spindles (Rossi et al., 2010). These results indicated that tip myonuclei expressed a different myosin program compared to other myonuclei in chicken foetal muscles.

We conclude that tip myonuclei of chicken limb foetal muscles display a molecular signature associated with a “signal transduction” identity, and a selective myosin program and contraction.

### Mononucleated *MYOD1*+ cells are heterogeneous with respect to cell cycle states

Myonuclei of foetal muscles are heterogeneous and display different molecular programs according to tip and central muscle domains (**Figs. 1**–**5**). We next addressed if mononucleated muscle cells also exhibited a regionalisation according to tips and central domains in foetal muscles. To address the heterogeneity of mononucleated cells, we implemented the scRNAseq dataset (6711 cells) of E10 limb muscle (Esteves de Lima et al., 2022) to our analysis (**Figs. 6, S3, S8**). Both snRNAseq and scRNAseq datasets of E10 limb muscle nuclei/cells illustrated myogenic progression from progenitors to differentiated mononucleated cells, rather than spatial heterogeneity **(Figs. 1, S3).** The increase of resolution from 0.2 (**Fig. S3, Table S3)** to 0.4 (**Fig. S8, Table S4**) of scRNAseq datasets led to two progenitor clusters (clusters 0,5) with *PAX7* and *PITX2* as respective markers of clusters 0 and 5), and three *MYOD1* clusters (clusters 1,2,3) (**Fig. 6**). *PAX7*, *PITX2* and *MYOD1* were described as being expressed uniformly in muscle progenitors and myoblasts of developing limbs (Esteves de Lima et al., 2022; L’Honoré et al., 2007). In *MYOD1* myoblast clusters, we noticed a spatial segregation according to cell cycle phases in both snRNAseq and scRNAseq datasets, one part being nuclei/cells in S/G2M phases and the other part being nuclei/cells in G1 phase (**Fig. 6A-D**). The expression of *KI67* associated with the G2/M phases (Uxa et al., 2021) and that of *CDKN1B* associated with the G1 phase (Bencivenga et al., 2021) were also heterogeneous in *MYOD1* clusters of both datasets (**Fig. S9A-D**). In the scRNAseq dataset at res 0.4, the three *MYOD1* clusters displayed different cell cycle states; one *MYOD1* cluster (cluster 1) was enriched in cells in G2M phases and *KI67* expression, one *MYOD1* cluster (cluster 2) was enriched in cells in G1 phase and *CDKN1B* expression (cluster 3) and another *MYOD1* cluster displayed no specific cell cycle feature (cluster 1) (**Figs. 6C,D, S8C, S9B,D**). To order the three *MYOD1* clusters according to myogenic progression, we performed stream analysis with the scRNAseq dataset of foetal muscle cells. As expected, the trajectory showed a direction from *PAX7* muscle progenitor clusters (clusters 0,5) to the muscle differentiated cluster (cluster 4) (**Figs. 6E, S9E**), illustrated with the successive and overlapping expression of *PAX7, MYOD, MYOG, TMEM8C* and *MYH1D* genes according to myogenic progression (**Fig. 6G**). Trajectory analyses showed a direction from *MYOD1* proliferating cluster 2 to *CDKN1B* post-mitotic cluster 3 (**Figs. 6E,F, S9E**), consistent with the cell cycle exit as being a key step of muscle differentiation onset during myogenic progression. In addition to the linear trajectory from progenitor to differentiation, we noticed a branch of the trajectory with a dead end in the *MYOD1* cluster 1 **(Fig. 6E,F** arrows**, Fig. S9E**), pointing to a bifurcation point in myogenic progression in between the proliferation step and the cell cycle exit step.

**Figure 6.**
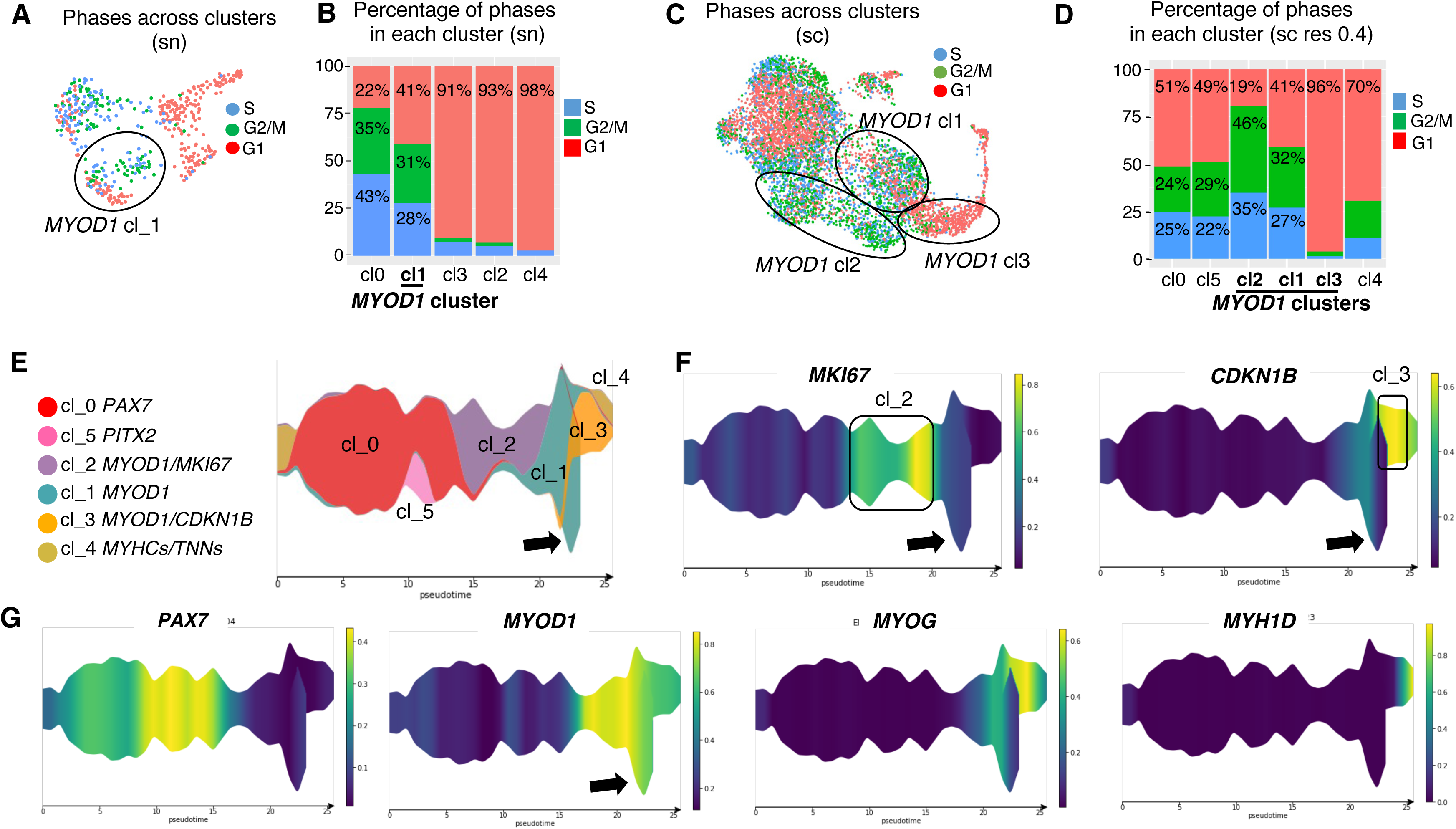
Cell cycle states in *MYOD*+ nuclei in E10 muscle snRNAseq dataset. (**A**) UMAP plot showing the distribution of the cell cycle phases across muscle nuclei of the E10 muscle snRNAseq dataset. (**B**) Stacked histogram showing the proportion of the cell cycle phases in each cluster of muscle snRNAseq datasets at resolution 0.5. (**C**) UMAP plot showing the distribution of the cell cycle phases across muscle cells of E10 muscle scRNAseq dataset. The three *MYOD1* clusters at resolution 0.4 are delineated. (**D**) Proportion of the cell cycle phases in each of the five clusters of E10 muscle scRNAseq datasets at resolution 0.4. (**E**) Dendrogram of the STREAM-derived inferred trajectory for the E10 muscle scRNAseq dataset showing the distribution of muscle clusters (resolution 0.4), ordering the MYOD1 clusters, cl2, 1 and 3. (**F**) Dendrogram plotting gene expressions levels for *MKI67* and *CDKN1B* genes on the inferred trajectory of the E10 muscle scRNAseq dataset, showing an enrichment of *MKI67* in *MYOD1-*cluster2 and *CDKN1B* in *MYOD1*-cluster 3. (**G**) Dendrogram plotting gene expressions levels for *PAX7, MYOD1, MYOG* and *MYH1D* genes on the inferred trajectory of the muscle scRNAseq dataset, showing the successive expression of muscle markers, highlighting the progression of myogenesis. Yellow indicates high levels of gene expression.

Despite a cell cycle regression step in our bioinformatics analysis, cells/nuclei within the *MYOD1* clusters segregate according to their cell-cycle status in both snRNAseq and scRNAseq datasets and according to myogenic progression

### Mononucleated *MYOD1*+ cells are regionalised according to tip and central domains

We then looked for markers of the three *MYOD1* clusters other than proliferative markers (**Fig. 7**). *FGFR4* (FGF receptor 4) was enriched in the *MYOD1* proliferative cluster (cluster 2) and the transcription factor *SOX8* was the top marker of cluster 1 (**Fig. 7A**), consistent with their known function in the first steps of the myogenic program (Bonnet et al., 2010; Marics et al., 2002; Schmidt et al., 2003). The fusion gene *TMEM8C* was a top marker of the *MYOD1* cluster more advance in the myogenic program (cluster 3) (**Fig. 7A, Table S2**). As for location, we know that *TMEM8C* transcripts are centrally located in limb foetal muscles (**Fig. 5A,B**) (Esteves de Lima et al., 2022). We looked for the location of *FGFR4* and *SOX8* transcripts in chicken limb muscles. FISH experiments to limb muscle sections at foetal stages with FGFR4 and SOX8 probes (**Fig. 7B-I**) showed that *FGFR4* and *SOX8* transcripts were located outside myosin+cells (**Fig. 7C,G,I**), consistent with their position in the myogenic program (Bonnet et al., 2010) (Schmidt et al., 2003) (Kim et al., 2022). Strikingly, *FGFR4* and *SOX8* transcripts were enriched in mononucleated muscle cells located close to tip myonuclei at the muscle/tendon interface of limb muscles at foetal stage (**Fig. 7B-I**). These results highlight a regionalisation of mononucleated muscle cells progressing from the tip domains (immature state) to the central domains (myoblasts that are engaged in the muscle differentiation program).

**Figure 7.**
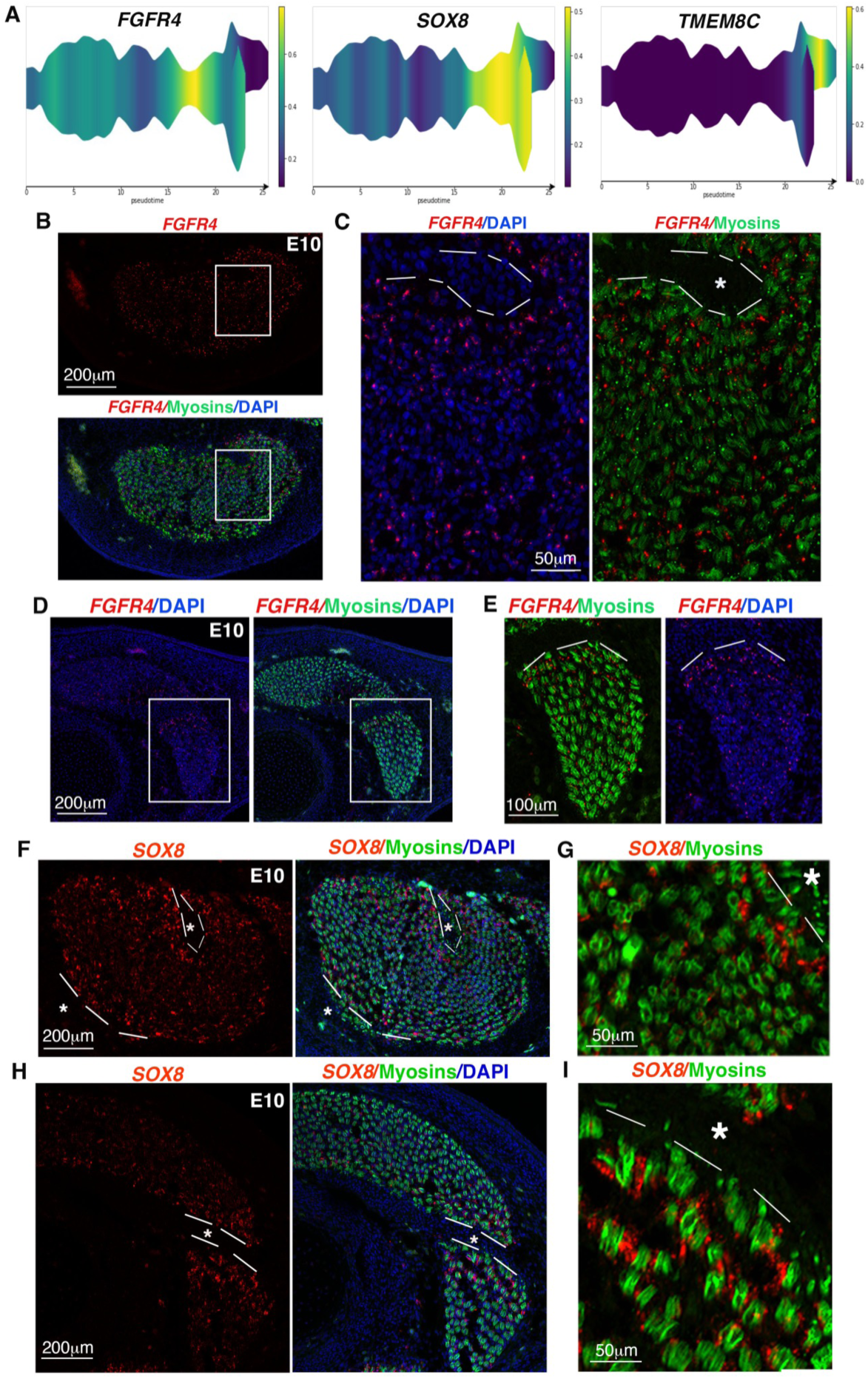
FGFR4 and *SOX8* expression is enriched in mononucleated myogenic cells close to tip myonuclei. (A) Dendrogram plotting gene expressions levels for *FGFR4*, *SOX8* and *TMEM8C* on the inferred trajectory of the E10 muscle scRNAseq dataset. Yellow indicates high levels of gene expression. (**B,C)** Fluorescent in situ hybridization to transverse limb sections focusing on the FCU (B,C) muscles with *FGFR4* probe (red) followed with an immunohistochemistry with MF20 antibody recognising myosins (green) and DAPI staining (blue). (C) High magnification squared in (B). (**D,E**) Fluorescent in situ hybridization to transverse sections of the anterior limb muscle (EMR) with *FGFR4* probe (red) followed with an immunohistochemistry with MF20 antibody recognising myosins (green) and DAPI staining (blue). (E) High magnification squared in (D). (**F**) Fluorescent in situ hybridization to transverse sections of the FCU limb muscles with *SOX8* probe (red) followed with an immunohistochemistry with MF20 antibody recognising myosins (green) and DAPI staining (blue). (**G**) High magnification of tip region of the FCU. (**H**) Fluorescent in situ hybridization to transverse sections of the anterior limb muscle (EMR) with *SOX8* probe (red) followed with an immunohistochemistry with MF20 antibody recognising myosins (green) and DAPI staining (blue). (**I**) High magnification of tip region of the EMR. *FGFR4* (B-E) and *SOX8* (F-I) expression was not evenly distributed in limb muscles, and showed an enriched expression in mononucleated muscles cells close to muscle/tendon interfaces. White dashed lines delineate the muscle/tendon interfaces. White stars indicate tendon location.

In this MS, we highlighted a molecular heterogeneity of myonuclei and mononucleated cells, which segregate according to tip and central muscle domains in foetal muscles of chicken limbs.

## Discussion

In this work we identified a molecular heterogeneity of foetal muscle nuclei, at the level of myonuclei but also at the level of mononucleated cells, according to tip and central muscle domains in developing chicken limbs.

### Myonuclei display either a tip or a central identity at early foetal stage

Using snRNAseq strategy combined with *in situ* hybridization experiments, we showed that foetal myonuclei can be divided into central and tip myonuclei in a complementary and non-overlapping manner, with distinct molecular signatures. Adult MTJ and MNJ markers were observed in the foetal tip and central domains, respectively, suggesting that the tip and central domains of foetal muscles prefigure the future adult MTJ and MNJ. Adult MTJ genes (Karlsen et al., 2023) found in tip foetal myonuclei include *COL22A1, ANKRD1, SMAD9, FGF6, MYH15.* The function of some of these genes has been described during development: SMAD9 as part of the BMP signalling is involved in fibroblast recruitment at muscle tips (Esteves de Lima et al., 2021) and FGF in the maintenance of tendon identity (Edom-Vovard et al., 2002). It is tempting to speculate that these foetal tip genes would have a similar function in adult MTJ. The central muscle domain concentrates nuclei expressing genes coding for post-synaptic components of NMJ but also for the master gene for myoblast fusion pointing to a close link between NMJ formation and myoblast fusion during development. Consistently, *COL25A1,* a top marker gene of central myonuclei of foetal muscles, has been shown to be involved in both processes, intramuscular innervation (Munezane et al., 2019; Tanaka et al., 2014) and myoblast fusion (Gonçalves et al., 2019) during mouse development.

In contrast to adult mouse and human muscles, where specialized MTJ and NMJ myonuclei correspond to less than 10% of myonuclei, reviewed in (Korb et al., 2024), at E10, central (*COL25A1*) and tip (*COL22A1*) markers are expressed in all myonuclei in a complementary manner, leaving no un-labelled myonucleus. However, at E14, the MTJ (*CHRNA1*+) and NMJ (*COL22A1*+) specialized domains have narrowed, while the central fusion domain (*TMEM8C+*) has not narrowed and continues to abut the tip domain. This leads to the emergence of myonuclei that are neither MTJ nor NMJ myonuclei. They could correspond to body myonuclei that would appear by *TMEM8C*+ myoblast accretion between the MTJ and NMJ domains during late foetal myogenesis.

### Tip foetal myonuclei as a signalling centre interacting with adjacent tendon fibroblasts

We identified a robust “signal transduction” transcriptional signature for tip myonuclei that was not observed in central myonuclei in limb foetal muscles of chicken embryos. This signature includes secreted molecules as well as receptor and effector genes, meaning that tip myonuclei both send and receive signals. We already know the function of several transduction pathways in the formation of the muscle/tendon interface during development. An FGF4 signal sent by tip myonuclei (Edom-Vovard and Duprez, 2004) maintains tendon identity by regulating tendon gene expression during limb and axial development (Brent et al., 2003; Brent and Tabin, 2004). BMP signalling was another signalling, which came up in the tip myonuclei molecular signature with *SMAD9* and *ID2* expressed by tip myonuclei, as a receiving signalling. A BMP signal sent by tendon fibroblasts (Wang et al., 2010) to tip myonuclei is involved in fibroblast recruitment at the tips of muscle fibres to generate a boundary of fibroblast-derived myonuclei at the MTJ (Esteves de Lima et al., 2021). In addition, we spotted the enrichment of genes involved in migration and cell-cell contact, previously characterized in the neural context (such as *VIP, ERBB4, NRCAM, DCX*). Along the same line, CXCR7, recently renamed as ACKR3 (Atypical ChemoKine Receptor3), is also involved in muscle stem cell migration during muscle regeneration (Kowalski et al., 2017). This enrichment of genes in migration and cell-cell contact in different contexts is consistent with the recognition step between muscle tips and tendon fibroblasts occurring at this developmental stage leading to muscle/tendon attachment. The molecular signature of tip myonuclei highlights reciprocal molecular interactions between tip myonuclei and tendon fibroblasts involved in the formation of the muscle/tendon interface during foetal development (**Fig. 8**).

**Figure 8.**
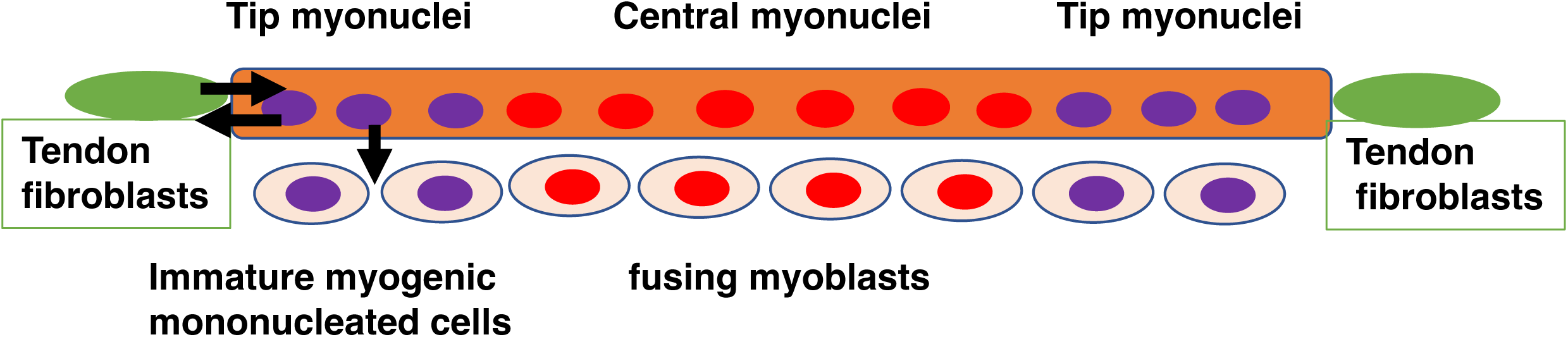
Regionalisation of myonuclei and mononucleated myogenic cells according to muscle domains at foetal stage. Schematic representation of tip (purple) and central (red) myonuclei in a myotube with tendon fibroblasts (green cells) at both extremities, with adjacent mononucleated myogenic cells at the tip domains (purple nuclei) and the central domain (red nuclei). Black arrows illustrate molecular interactions between tip myonuclei, and adjacent tendon fibroblasts and adjacent immature mononucleated myogenic cells.

### Tip foetal myonuclei as a signalling centre interacting with adjacent mononucleated muscle cells to regulate myogenesis

In addition to reciprocal molecular interactions between muscle tips and tendon fibroblasts, we believe that there are molecular interactions between multinucleated muscle fibres and mononucleated muscle cells at the tip domain. In this study, we identified an enrichment of mononucleated muscle cells expressing the FGF receptor FGFR4 at the tip domain, while the nearby tip myonuclei produce FGF ligands (FGF6, FGF4). These tip mononucleated muscle cells express both *FGFR4* and *SOX8* genes that are described as progenitor markers upstream of MYOD1 in a fibroblast-to-myogenic conversion culture system (Kim et al., 2022). *FGFR4* is recognized to act downstream of *PAX3* and upstream of *MYOD* during chicken and mouse development (Bonnet et al., 2010; Lagha et al., 2008). Both *FGFR4* and *SOX8* genes have been described as markers of satellite cells that prevent muscle differentiation of satellite cells (Schmidt et al., 2003; Yeh et al., 2025). An attractive hypothesis is that the FGFs produced by tip myonuclei would signal to the adjacent mononucleated muscle cells expressing the FGF receptor FGFR4 to regulate their entry into myogenesis, maintaining them in an immature state waiting to fuse. These pausing muscle cells would correspond to the SOX8-associated branch of the inferred trajectory, consistent with SOX8 function inhibiting myogenic progression (Schmidt et al., 2003). Signalling from tip myonuclei would thus regulate the size of the pool of mononucleated muscle cells, consistent with the role of FGF6 and FGFR4 in regulating the size of muscle stem cell pool in postnatal mice (Yeh et al., 2025; Zofkie et al., 2021). The tip domain would be a region enriched with pausing mononucleated muscle cells, while the central domain would concentrate fusing myoblasts, indicating a myogenic progression from tip (immature) to middle (fusion) in foetal muscles, under the regulation by tip myonuclei (**Fig. 8**).

## Conclusion

We identified a regionalisation of the transcriptional signatures of myonuclei and mononucleated myogenic according to tip and central domains in foetal limb muscles of developing chicken embryos. Foetal myogenesis corresponds to a time of intricated processes such muscle/tendon interface formation and muscle growth. We propose that foetal myonuclei at the tip domain act as a signalling centre involved in muscle/tendon recognition and attachment but also in muscle growth by regulating myogenesis.

## Materials and Methods

### Chicken embryos

Fertilized chicken eggs (JA57 strain) from commercial source (EARL Les Bruyères, Dangers, France) were incubated at 37.5°C in a humidified incubator until appropriate stages. Embryos were staged according to days of *in ovo* development. All experiments with chicken embryos were performed before E14 and consequently are not submitted to a licensing committee, in accordance with European guidelines and regulations.

### Seurat clustering analysis

The sample collection, scRNAseq protocol and clustering analysis for E10 whole limb cells are already published in (Esteves de Lima et al., 2021).

#### Sample preparation for the single-nucleus E10 dataset

Two forelimbs (fingers excluded) from one E10 embryo were dissected in PBS and snap-frozen in liquid nitrogen. The subsequent steps were performed at 4°C. The tissue was transfered in lysis buffer (10mM Tris-HCl, 10mM NaCl, 3mM MgCl2, 0.1% Nonidet-TM P40 in Nuclease-Free Water), homogenized and coarsly dissociated with a pipette. Nuclei were extracted by a first round of Dounce homogenizer (20 strokes of loose pestle), followed by a 10 min incubation with gentle mixing and a second round of Dounce homogenizer (20 strokes of tight pestle). Cell debris were removed by a 70 μm filtration followed by a 40 μm filtration, applying wash buffer (PBS, 2% BSA, 0,2U/μl RNase inhibitor (Roche)) after each filtration. To label nuclei, the nuclear suspension was centrifuged at 500 g for 8 min and resuspended in PBS/2% BSA and centrifuged again at 500 g for 8 min, resuspended and incubated in wash buffer/10 μg/ml DAPI in the dark for 10 min. After adding PBS/2% BSA, the nuclear suspension was centrifuged at 500 g for 8 min, resuspended and filtrated on a 30 μm filter with wash buffer. For quality control and counting, the DAPI-labelled nuclear suspension was then run on the Genom’IC platform FACS, Institut Cochin, Paris.

10 000 nuclei were loaded into the 10x Chromium Chip with the Single Cell 3′ Reagent Kit v3 according to the manufacturer’s protocol. Libraries were then sequenced by pair with a HighOutput flowcel using an Illumina Nextseq 500 with the following mode (150 HO): 28 base-pairs (bp) (Read1), 56 bp (Read 2) and 8 bp (i7 Index). An average of 50 000 reads per cell were sequenced and analysed with Cell Ranger Single Cell Software Suite 4.0.0 by 10x Genomics. Raw base call files from the Nextseq 500 were demultiplexed with the cellranger mkfastq pipeline into library-specific FASTQ files. The FASTQ files for each library were then processed independently with the cellranger count pipeline. This pipeline used STAR21 to align cDNA reads to the Gallus gallus genome (Sequence: GRCg6a, Gallus gallus pre-mRNA reference). To remove ambient RNA contamination, a corrected count matrix was generated from the cellranger count matrices using SoupX (default parameters) (Young 2020: 10.1093/gigascience/giaa151) under R (v4.0.1) (Team, R.C. (2018). R: A language and environment for statistical computing. (R Foundation for Statistical Computing, Vienna, Austria.)).

#### Seurat clustering of whole limb E10 snRNAseq dataset

The Seurat package (v4) (Stuart et al., 2019) under R (v4.0.1) (Team, R.C. (2018). R: A language and environment for statistical computing. (R Foundation for Statistical Computing, Vienna, Austria.)) was used to perform downstream clustering analysis on snRNAseq data (Macosko et al., 2015) (see **Table S5** for parameter and cut-off values). Nuclei went through a classical Quality Control using the number of detected genes per cell (nFeatures), the number of mRNA molecules per cell (nCounts) and the percentage of expression of mitochondrial genes (pMito) as cut-offs. Potential doublets were identified by running the Scrublet algorithm (Wolock et al., 2019) and then removed from the dataset. Outliers on a nFeature vs nCount plot were manually identified and removed from the dataset. Gene counts for cells that passed the above selections were normalized to the total expression and Log-transformed with the NormalizeData function of Seurat (scale factor = 10 000). Highly variable genes were detected with the FindVariableFeatures function (default parameters). Cell cycle effect was regressed out using the ScaleData function. Using highly variable genes as input, principal component analysis was performed on the scaled data in order to reduce dimensionality. Statistically significant principal components were determined by using the JackStrawPlot and the ElbowPlot functions. Cell clusters were generated with the FindNeighbors/FindClusters functions (default parameters except for the number of selected PCs). Different clustering results were generated at different resolutions and for different sets of PCs. Non-linear dimensional reduction (UMAP) and clustering trees using Clustree (Zappia and Oshlack, 2018) were used to visualize clustering results and select for the most robust and relevant result. Markers for each cluster were found using the FindAllMarkers function of Seurat (using highly variable genes as an input, default parameters otherwise) that ran Wilcoxon rank sum tests (p-val adjusted < 0.05). The clusters identified as Muscle clusters by the differential expression of classical muscle markers (*PAX7, MYOD1, MYOG, MYH1D*) represented the minority of limb cells, in addition to other clusters encompassing the expected cell populations present in developing limb tissues such as connective tissues, vessels/blood, neural crest cells and ectoderm (**Fig. S1)** as in scRNAseq datasets of limb cells (Esteves de Lima et al., 2021).

#### Seurat clustering of Muscle subsets

For the sc-and sn-RNAseq datasets, we selected the Muscle (M) clusters and extracted them from the post-Quality Control object to generate a M subset of the whole limb dataset. The sizes of M datasets were 6711 cells (M-E10-sc) and 507 nuclei (M-E10-sn). M datasets were then processed again through the pipeline described above, from the Normalization to clustering and marker identification steps. In the snRNAseq dataset, to identify highly specific markers for each cluster, the marker list of M-E10-sn (**Table S1**) was further filtered (pct.1> 0.40; pct.2<0.25) to generate the filtered marker lists (**Table S2**), pct.1 being the fraction of cells in the cluster of interest expressing the marker gene of interest and pct.2, the fraction of cells in the other clusters expressing that marker gene. The M datasets were analyzed using classical Seurat tools such as Feature plots, Violin plots, Dimplots, Heatmaps and Dotplots. Gene expression was defined by ‘gene log-normalized count>0’. The R package ggplot2 v3.5.2 (Wickham, 2016) was used to generate custom feature plots highlighting gene co-expression. Population intersection plots were generated with the R package UpSetR v1.4.0 (Conway et al., 2017).

### STREAM trajectory inference

The STREAM pipeline (v1.0) (Chen et al., 2019) under Python (v3.7.9) was used to perform trajectory inference analysis on the M-E10-sc dataset (see Table S1 for parameter values). The M-E10-sc dataset was imported into the STREAM environment under the h5ad format to build the trajectory on the Seurat UMAP dimension reduction. To perform gene expression analysis on top of trajectory inference, the M-E10-sc dataset was also imported under the 10X format and we generated an anndata object combining the expression matrix (RNA slot) of the M-E10-sc 10x object with the dimension reduction matrix of the M-E10-sc h5ad object. Trajectories were inferred with the elastic_principal_graph function, using the umap reduction. Trajectories were visualized either on a UMAP (plot_dimension_reduction function) or as a branching tree (plot_flat_tree function) or as a dendrogram (plot_stream function), color-coded with the Seurat cluster labels or gene expression levels.

### Reactome analyses of the cluster-associated markers

GO analysis was performed on the cluster 4 filtered marker list using the ‘Analyze Gene List’ tool from the Reactome Pathway Database (version 97) (Fabregat et al., 2017; Wu and Haw, 2017) (**Table S6**). GO terms were selected based on their p-value (< 0.05) and plotted as a dotplot, highlighting their p-value and gene number.

### Comparison of adult human MTJ and foetal chicken tip myonuclei snRNAseq datasets

#### Analysis of human snRNAseq dataset

Publicly available snRNA-seq data from human semitendinosus muscle–tendon tissue (three healthy male donors; ArrayExpress E-MTAB-12529; (Karlsen et al., 2023)) were reanalysed using Seurat (v5.4.0). Raw count matrices from three samples (S1, S3 and S4) were individually filtered (>200 genes and >500 UMIs), corrected for ambient RNA using SoupX (v1.6.2), and subjected to doublet removal using Scrublet (v0.2.3). Nuclei with fewer than 300 detected genes after correction were excluded. Samples were normalised using SCTransform and integrated using the Seurat SCT workflow (2,000 features). PCA, UMAP and clustering were performed using 30 PCs (resolution 0.5). Myonuclei were identified by *TTN* expression and the absence of expression of markers associated with non-myonuclear cell populations (*PAX7, COL1A2, COL3A1, FAP, DCN, TCF7L2, PDGFRA, COMP, MKX, ACTA2, MYH11, RGS5, VWF, PECAM1, CD163,* and *F13A1*). Extracted myonuclei (n = 20,614) were reanalysed using SCTransform and clustered using 20 PCs (resolution 0.1), yielding five clusters annotated by *MYH7* (slow), *MYH1*/*MYH2* (fast) and *COL22A1*/*NCAM1*/*SORBS2* (MTJ; cluster 3). MTJ-enriched genes were identified using Seurat FindConservedMarkers, requiring Bonferroni-adjusted *P* < 0.05 and average log2 fold change > 0.35 in all three samples, and expression in >10% of MTJ myonuclei in at least one sample, yielding 354 genes.

#### Chicken MTJ-enriched genes and cross-species comparison

Chicken MTJ-enriched genes were obtained from the E10 snRNA-seq analysis described above (Bonferroni-adjusted *P* < 0.05; average log2 fold change > 0.35; n = 332). Of these, 318 genes had annotated symbols and were mapped to 294 unique human orthologues using Ensembl BioMart (biomaRt v2.62.1; Ensembl release 115; accessed on 6 May 2026), including five genes matched by identical symbols. Comparison with the 354 human MTJ-enriched genes identified 41 shared genes, 313 human-only genes and 253 genes unique to the chicken-derived orthologue set (**Fig. S6A**; ggVennDiagram v1.5.7). Shared gene expression was visualised using Seurat DotPlot (**Fig. S6B**), myonuclear populations using UMAP and canonical marker expression (**Fig. S6C**), and selected genes (*COL22A1*, *SMAD9*, *FGF6* and *MYH15*) using FeaturePlot and VlnPlot (**Fig. S6D**).

### In situ hybridization

<u>Colorimetric or Fluorescent in situ hybridization (FISH)</u> was performed on tissue sections as described (Hirsinger et al., 2024). Chicken embryos were fixed in 4% paraformaldehyde (Sigma-Aldrich) overnight at 4°C, then processed in 7.5%/15% gelatin/sucrose (Sigma-Aldrich) for cryostat sections. Limbs were cut in 12-μm transverse or longitudinal sections and processed for *in situ* hybridization. Alternating serial sections were hybridized with probe 1, probe 2 and probe 3 to allow comparison of expression domains on adjacent sections of the same limb. Digoxigenin-labelled or Fluorescein-labelled mRNA probes were prepared as described for TMEM8C (Esteves de Lima et al., 2022), COL22A1 (Nicolas et al., 2024), FGFR4 (Bonnet et al., 2010), COL12A1 (Lejard et al., 2011), ETV5/ERM (Brent and Tabin, 2004). The primers used for the following PCR probes, CHRNA1, COL24A1, COL25A1, CXCR7, DCLK3, ERBB4, MHY15, SMAD9, SMTNL2, SOX8, UACA and VIP, are listed in Table S8. Colorimetric in situ hybridization were revealed with an antibody against Digoxigenin labelled with alkaline phosphatase. FISH were revealed with antibodies (against Digoxigenin or Fluorescein) labelled with POD. Fluorescence was revealed with the TSA+ Cyanin 3 or 5 system (AKOYA biosciences) (CHRNA1, COL24A1, COL25A1, CXCR7, DCLK3, ERBB4, MHY15, SMAD9, SMTNL2, SOX8, UACA and VIP).

<u>In situ hybridization chain reaction (HCR-ISH)</u> was performed on tissue sections using the HCR v3.0 protocol for tissue sections (Molecular Instruments), with custom-designed DNA probe sets. Tissue sections were pre-hybridized in 30% probe hybridization buffer for 10 min at 37°C in a humidified chamber. Hybridization was performed overnight (12–16 h) at 37°C with 0.4 pmol of each probe mixture in 100 µL of probe hybridization buffer. Following hybridization, sections were washed sequentially at 37°C for 15 min each in 75%, 50%, and 25% probe wash buffer diluted with 5× SSCT, followed by 100% 5× SSCT. Sections were then washed in 5× SSCT for 5 min at room temperature. For signal amplification, sections were pre-incubated in amplification buffer for 30 min at room temperature. Fluorescently labelled DNA hairpins (6 pmol each) were snap-cooled at 95°C for 90 s and allowed to cool to room temperature for 30 min before being added to 100 µL of amplification buffer. Sections were incubated with the hairpin solution overnight (12–16 h) at room temperature in the dark. Excess hairpins were removed by washing twice for 30 min and once for 5 min in 5× SSCT at room temperature. The primers used for HCR-ISH, CHRNA1, COL22A1, COL25A1 and TMEM8C, are listed in Table S9.

### Immunohistochemistry

Fluorescent and HCR *In situ* hybridization to limb sections or to cells was followed with immunohistochemistry using the MF20 monoclonal antibody (DSHB cat. # MF20, undiluted supernatant), which recognized sarcomeric myosin heavy chains. The MF20 monoclonal antibody developed by D.A. Fischman and A. Kawakami, respectively were obtained from the Developmental Studies Hybridoma Bank developed under the auspices of the NICHD and maintained by the University of Iowa, Department of Biology Iowa City, IA 52242, USA. Secondary antibodies were conjugated with Alexa 488, Alexa 555 or Alexa 647 (Invitrogen). Nuclei were visualized with DAPI (Sigma-Aldrich) staining.

### Image capture

After *in situ* hybridization and/or immunohistochemistry experiments images were obtained using an Apotome.2 epifluorescence microscope (Zeiss).

## Acknowledgements

We thank the InforBio (ex ARTbio) platform of the IBPS for support in bioinformatics analyses. Sequencing and genome mapping of the snRNAseq and scRNAseq datasets were performed by the GENOM’IC platform at the Cochin Institute (Paris).

## Ethics approval and consent to participate

All experiments with chicken embryos were performed before E14 (embryonic Day 14) of development) and consequently are not submitted to a licensing committee, in accordance with European guidelines and regulations.

## Author contribution

DD conceived the project. TK, CB and MAB performed the experiments while TG and EH conducted the bioinformatics analyses. TK, EH and DD analyzed the data. DD wrote the original draft. TK, MCD, EH and DD reviewed and edited the original draft. DD supervised the project and acquired funding.

## Data and Ressource availability

All data are available within the Article and Supplementary Files. Source data for all graphics are provided with this paper. The scRNAseq dataset have been deposited in the NCBI Gene Expression Omnibus database (https://www.ncbi.nlm.nih.gov/geo/), and is accessible under the number GSE166981. The snRNAseq analysis was performed with standard pipelines, as described in the Methods. Scripts will be made available upon request. Further information and requests for resources and reagents will be fulfilled upon request. Microscopy data reported in this paper will be shared upon request.

## Supplementary Figure legends

**Figure S1**

**Single-nucleus RNA-sequencing datasets of whole limb nuclei from E10 chicken embryos**

(**A**) UMAP plot showing the distribution of clusters (3174 nuclei) in E10 whole limb snRNAseq dataset. Muscle clusters are circled in red, while connective tissue clusters are circled in black.

(**B**) Stacked histogram showing the percentage of nuclei per cluster in E10 whole limb snRNAseq datasets. Muscle clusters 6,7 and 10 are boxed in red.

**(C)** Violin plots showing the log-normalized expression levels of known markers across clusters in the E10 whole limb snRNAseq dataset, allowing for the identification of the ectoderm clusters (9,12) with *WNT6*, the connective tissue clusters (2,3,0,4,5) with *PDGFRA* and *PRRX1*, the tendon clusters (1,8) with *MKX* and *SCX*, the perichondrium cluster 15 with *SATB2*, the neural crest cluster 14 with *SOX10*, the smooth muscle cluster 13 with *ACTA2* and the endothelial cells cluster 11 with *KDR*.

(**D**) Violin plots showing the log-normalized expression levels of *PAX7*, *MYOD*, *MYH1D* genes across clusters, in E10 whole limb snRNAseq datasets, allowing for the identification of the muscle clusters 6,7,10.

**Figure S2**

**Clustering analysis of the muscle subset of the E10 whole limb snRNAseq dataset**

(**A**) Dotplot showing the average expression levels and the fraction of expressing-nuclei for the top10 markers (ordered by fold-change) of each of the five clusters in the E10 muscle snRNAseq dataset, ordered according to myogenic progression.

(**B**) Feature plots showing the log-normalized expression levels across clusters for the following muscle markers, *PAX3*, *FGFR4*, *MYOD1*, *MYOG*, *TMEM8C*, *MYH1B, MYH1G* and *MYH7B* in the E10 muscle snRNAseq datasets Corresponding violin plots are shown in Fig. 2C.

(**C**) Feature and violin plots showing the distribution and log-normalized expression levels for the muscle differentiation genes, *TNNT1, TNNT2* and *DES* across nuclei and clusters.

**Figure S3**

**Myogenic progression illustrated by the analysis of the muscle subset of the E10 whole limb single-cell RNA-sequencing dataset (resolution 0.2)**

(**A**) UMAP plot showing the distribution of the 3 muscle clusters.

(**B**) Stacked histogram showing the percentage of cells per muscle cluster.

(**C**) Stacked histogram showing the percentage of nuclei per cluster for the three first clusters according to myogenic progression (clusters 0, 1 and 3) of the muscle snRNAseq dataset.

(**D**) Violin and feature plots showing the log-normalized expression levels and distribution for the following muscle markers, *PAX7, MYOD1, MYOG* and *MYH1D* across clusters and cells in the E10 muscle scRNAseq dataset at resolution 0.2.

(**E**) Heatmap showing the relative expression of the top 10 markers across all cells of the three clusters (in the muscle scRNAseq dataset at resolution 0.2), ordered according to myogenic progression. Upregulated genes are in yellow, while downregulated genes are in purple.

(**F**) Feature plots with high magnifications showing the log-normalized expression levels for the fibroblast markers *PDGFRA, HIC1* and *TWIST2* across cells pointing to the dual fibroblast/muscle cells.

**Figure S4**

**COL22A1 is a reliable marker for tip myonuclei in E10 muscle single-nucleus RNA-sequencing dataset and limb foetal muscles**

(**A**) UMAP plots at different clustering resolutions, 0.1, 0.2, 0.4, 0.5 and 0.9, showing the evolution of the distribution of muscle clusters in the E10 muscle snRNAseq dataset of chicken limb muscles at E10.

(**B**) Clustree tree showing the bifurcations across resolution 0.2, 0.4 and 0.9. The tip myonuclei cluster appears at resolution 0.4 and is observed until resolution 1.

(**C,D**) Feature plots showing the distribution of *COL22A1*+nuclei across clusters in E10 limb muscle (C) and E10 whole limb (D) snRNAseq datasets.

(**E-J**) Colorimetric in situ hybridization to longitudinal (E-H) and transverse (I,J) limb sections for *COL22A1* (blue) followed by an immunostaining with the MF20 antibody to visualize myosins (brown) from E7, E9 and E12 chicken embryos. (F,H,J) are high magnifications of muscles squared in (E,G,I), respectively.

**Figure S5**

**Molecular markers of tip myonuclei**

(**A,B**) Fluorescent in situ hybridization to transverse limb muscle sections of E10 chicken embryos followed with an immunohistochemistry with the MF20 antibody recognising myosins (green) and DAPI staining (blue) with *DCLK3* probe (red). (A) Low magnification of transverse limb section with *DCLK3*/DAPI staining. (B) is a focus of the FCU muscle (ventral and posterior muscle) with *DCLK3/*DAPI staining and *DCLK3*/myosins staining. Dashed lines indicate the muscle/tendon interface.

**(C,D**) Fluorescent in situ hybridization to adjacent transverse limb muscle sections of E10 chicken embryos followed with an immunohistochemistry with the MF20 antibody recognising myosins (green) and DAPI staining (blue) with *DCX* (C) and *COL12A1* (D) probes (red). (C’,D’) are high magnifications of part of the dorsal muscle squared in upper panels. Dashed lines indicate the muscle/tendon interface. Arrows point to *DCX* expression in tip myonuclei, while *COL12A1* labels tendons.

(**E**) Violin and feature plots showing the log-normalized expression levels and the distribution of the *UACA* gene across clusters of E10 muscle snRNAseq dataset.

(**F**,**G,G’**) Fluorescent in situ hybridization to transverse limb muscle sections of E10 chicken embryos followed with an immunohistochemistry with the MF20 antibody recognising myosins (green) and DAPI staining (blue) with *UACA* probe (red). (F) Low magnification of transverse limb section with *UACA*/DAPI staining showing the regionalised *UACA* expression in limb muscles. (G) is a focus of a dorsal and posterior muscle showing *UACA* expression in tip myonuclei. Dashed lines indicate the muscle/tendon interface. (G’) is a high magnification of the muscle shown in (G) focusing on muscle tip.

(**H**) UpSet plot showing nuclei numbers per gene combination between the top 6 genes of cluster 4 filtered marker list (*COL22A1, ERRB4, DCLK3 COL24A1, UACA, VIP*). The size of each population expressing a gene of interest is shown at the bottom left of the plots.

**Figure S6**

**Comparison of the molecular signatures between chicken tip foetal myonuclei with human adult MTJ myonuclei**

(**A**) Venn diagram of MTJ myonuclei genes from human adult muscle snRNAseq datasets (Karlsen et al., 2023) and chicken E10 muscle snRNAseq dataset.

(**B**) Dotplot showing the average expression levels and the fraction of expressing-nuclei for 41 shared genes between the human adult muscle snRNAseq datasets (Karlsen et al., 2023) and the chicken E10 muscle snRNAseq dataset.

(**C**) UMAP plot showing the myonuclei distribution from the three human samples (Karlsen et al., 2023) and dotplot showing the average expression levels and the fraction of expressing-nuclei for the main markers in each of the five clusters of the human adult muscle snRNAseq datasets (Karlsen et al., 2023).

(**D**) Feature and Violin plots showing the distribution and the log-normalized expression levels of four shared genes, *COL22A1, SMAD9, FGF6* and *MYH15* across myonuclei and clusters, in human muscle datasets; genes for which the expression is validated in tip myonuclei of chicken E10 muscles.

**Figure S7**

**NMJ post-synaptic markers in E10 muscle snRNAseq dataset and in E10 chicken limb muscles**

(**A**) Feature and violin plots showing the distribution and log-normalized expression levels of the *VAV3, LRP4, CHRNG, CAMK4* post-synaptic genes across nuclei and clusters (genes present in non-filtered gene list of cluster 2 and listed by fold-changes high to low)

(**B**) Feature and violin plots showing the distribution and log-normalized expression levels of *CHRNA1* and *ETV5* post-synaptic genes across nuclei and clusters.

(**C**) Colorimetric in situ hybridization to longitudinal muscle sections of E10 chicken limbs for CHRNA1, ETV5 and FGF4 probes (blue), showing the central location of *CHRNA1, ETV5* transcripts and the tip location of *FGF4* transcripts.

**Figure S8**

**Clustering analysis of the E10 muscle scRNAseq dataset at resolution 0.4**

(**A**) UMAP plot showing the distribution of the six muscle clusters at E10, at resolution 0.4, and associated cell composition of each cluster: the two progenitor clusters (0,5), the three MYOD1 clusters (2,1,3) and the myosin cluster 4.

(**B**) Stacked histogram showing the percentage of cells per cluster from the E10 muscle scRNAseq dataset at resolution 0.4.

(**C**) Violin plots showing the log-normalized expression levels for the following muscle markers, *PAX7, PITX2, FGFR4, MYOD1, MYOG, TMEM8C, MKI67, SOX8, CDKN1B, TNNT1, TNNT3* and *MYH1D,* across clusters from the muscle scRNAseq dataset at resolution 0.4.

(**D**) Heatmap showing the relative expression of the top 10 markers across all cells of the six muscle clusters (from the muscle scRNAseq dataset resolution 0.4), ordered according to myogenic progression. Upregulated genes are in yellow, while downregulated genes are in purple.

**Figure S9**

**Cell cycle markers in the muscle snRNAseq and scRNAseq datasets and STREAM inferred trajectory in the muscle scRNAseq dataset**

(**A,B**) UMAP plot showing the distribution of MKI67+ nuclei/cells and CDKN1B+ nuclei/cells according to cell cycle phases across nuclei/cells in snRNAseq (A) and scRNA seq (B) E10 muscle datasets. *MKI67* is associated with S/G2/M phases, while *CDKN1B* is associated with G1 phase in both datasets.

(**C,D**) Feature plots showing the log-normalized expression levels of *MKI67* and *CDKN1B* across nuclei/cells in E10 muscle snRNAseq (C) and scRNA seq (D) datasets, showing the non-uniform distribution of *MKI67* and *CDKN1B* expression.

(E) UMAP and branching tree plot of the inferred trajectory performed on the scRNAseq muscle dataset (res 0.4), showing the distribution of muscle clusters.

## Supplementary Table legends

**Table S1**

**Marker list for the five muscle clusters of E10 limb muscle single-nucleus RNAseq dataset at resolution 0.5**

Excel file of the differentially-expressed genes in each cluster, ordered by decreasing fold-change, from snRNAseq datasets of E10 limb muscle nuclei. The ultimate_gene_names have been identified by combined data-mining of the Ensembl and NCBI Entrez databases. Pct.1 is the fraction of marker-expressing cells in the cluster of interest. Pct.2 is the fraction of marker-expressing cells in the other clusters. In the column “cluster” is listed the cluster the marker is associated with for a given row. In the column “cluster_list” are listed all the clusters this marker is associated with.

**Table S2**

**Filtered marker list for the five muscle clusters of E10 limb muscle single-nucleus RNAseq dataset at resolution 0.5**

Table S2 corresponds to Table S1 after filtration on ptc.1 and pct.2 values. Only genes with pct.1 > 0.40 and pct.2 < 0.25 were selected. Excel file of the differentially-expressed genes respecting the above thresholds in each cluster, ordered by decreasing fold-change, from snRNAseq dataset of E10 limb muscle nuclei. Key muscle genes and genes mentioned in this study were highlighted in the colour of in each cluster. Markers selected for ISH experiments were highlighted in bold. Genes for which the expression is known in developing muscles are also highlighted in bold and referenced in the text.

**Table S3**

**Marker list for the three muscle clusters of E10 limb muscle single-cell RNAseq dataset at resolution 0.2**

Excel file of the differentially-expressed genes in each cluster, ordered by decreasing fold-change from snRNAseq dataset of E10 limb muscle cells. The ultimate_gene_names have been identified by combined data-mining of the Ensembl and NCBI Entrez databases. Pct.1 is the fraction of marker-expressing cells in the cluster of interest. Pct.2 is the fraction of marker-expressing cells in the other clusters. In the column “cluster” is listed the cluster the marker is associated with for a given row. In the column “cluster list” are listed all the clusters this marker is associated with. Key muscle genes and genes mentioned in this study were highlighted in the colour of in each cluster. Genes for which the expression is known in developing muscles are highlighted in bold and referenced in the text.

**Table S4**

**Marker list for the six muscle clusters of E10 limb muscle single-cell RNAseq dataset at resolution 0.4**

Excel file of the differentially-expressed genes in each cluster, ordered by decreasing fold-change from snRNAseq datasets of E10 limb muscle cells. The ultimate_gene_names have been identified by combined data-mining of the Ensembl and NCBI Entrez databases. Pct.1 is the fraction of marker-expressing cells in the cluster of interest. Pct.2 is the fraction of marker-expressing cells in the other clusters. In the column “cluster” is listed the cluster the marker is associated with for a given row. In the column “cluster list” are listed all the clusters this marker is associated with. Key muscle genes and genes mentioned in this study were highlighted in the colour of in each cluster. Genes for which the expression is known in developing muscles are highlighted in bold and referenced in the text.

**Table S5**

**Values for the parameters and cut-offs used in the Seurat and the STREAM analyses**

**Table S6**

**Reactome analyses on the cluster 4-associated filtered marker list and the cluster 2-associated filtered marker list from the snRNAseq dataset at resolution 0.5**

Excel files of the enriched Pathway names with Entities pValue < 0.05, ordered by increasing Entities pValues. Only the exclusive markers associated with cluster 4 or cluster 2 were considered for the analysis.

For cluster 4 marker list, 66 out of the 94 markers were found in Reactome, where 464 pathways were hit by at least one of them.

For cluster 2 marker list, 53 out of the 83 markers were found in Reactome, where 270 pathways were hit by at least one of them.

Pathway names of higher order in the Reactome hierarchy with high #Entities found were highlighted in colours and plotted on Fig. 3A,B.

**Table S7**

**Reactome analysis on the cluster 4 marker list (versus cluster 2) and the cluster 2 marker list (versus cluster 4) from the snRNAseq dataset at resolution 0.5**

Exel file of the enriched Pathway names with Entities pValue < 0.05, ordered by increasing Entities pValues for the cluster 4 versus cluster 2 comparison. 74 out of the 117 markers were found in Reactome, where 522 pathways were hit by at least one of them.

Excel file of the marker list, enriched in cluster 4 versus cluster 2.

Excel file of the enriched marker lists in cluster 2 vs cluster 4 and Pathway names with Entities pValue < 0.05, ordered by increasing Entities pValues, and marker lists for cluster 2 vs cluster 4 comparison. 29 out of the 47 markers were found in Reactome, where 190 pathways were hit by at least one of them.

Excel file of the marker list, enriched in cluster 2 versus cluster 4.

**Table S8**

**Primer list for Fluorescent-ISH experiments**

**Table S9**

**Primer list for HCR-ISH experiments**

## References

Aittaleb, M., Martinez-Pena, Y. V. I. and Akaaboune, M. (2017). Spatial distribution and molecular dynamics of dystrophin glycoprotein components at the neuromuscular junction in vivo. J Cell Sci 130, 1752–1759.

Armand, A. S., Laziz, I. and Chanoine, C. (2006). FGF6 in myogenesis. Biochim Biophys Acta 1763, 773–778.

Ayanlaja, A. A., Xiong, Y., Gao, Y., Ji, G., Tang, C., Abdikani Abdullah, Z. and Gao, D. (2017). Distinct Features of Doublecortin as a Marker of Neuronal Migration and Its Implications in Cancer Cell Mobility. Front Mol Neurosci 10, 199.

Bencivenga, D., Stampone, E., Roberti, D., Della Ragione, F. and Borriello, A. (2021). p27(Kip1), an Intrinsically Unstructured Protein with Scaffold Properties. Cells 10.

Bonnet, A., Dai, F., Brand-Saberi, B. and Duprez, D. (2010). Vestigial-like 2 acts downstream of MyoD activation and is associated with skeletal muscle differentiation in chick myogenesis. Mech Dev 127, 120–136.

Brent, A. E., Schweitzer, R. and Tabin, C. J. (2003). A somitic compartment of tendon progenitors. Cell 113, 235–248.

Brent, A. E. and Tabin, C. J. (2004). FGF acts directly on the somitic tendon progenitors through the Ets transcription factors Pea3 and Erm to regulate scleraxis expression. Development 131, 3885–3896.

Charvet, B., Guiraud, A., Malbouyres, M., Zwolanek, D., Guillon, E., Bretaud, S., Monnot, C., Schulze, J., Bader, H. L., Allard, B. et al. (2013). Knockdown of col22a1 gene in zebrafish induces a muscular dystrophy by disruption of the myotendinous junction. Development 140, 4602–4613.

Chen, H., Albergante, L., Hsu, J. Y., Lareau, C. A., Lo Bosco, G., Guan, J., Zhou, S., Gorban, A. N., Bauer, D. E., Aryee, M. J. et al. (2019). Single-cell trajectories reconstruction, exploration and mapping of omics data with STREAM. Nat Commun 10, 1903.

Cochaud, S., Meunier, A. C., Monvoisin, A., Bensalma, S., Muller, J. M. and Chadéneau, C. (2015). Neuropeptides of the VIP family inhibit glioblastoma cell invasion. J Neurooncol 122, 63–73.

Comai, G. and Tajbakhsh, S. (2014). Molecular and cellular regulation of skeletal myogenesis. Curr Top Dev Biol 110, 1–73.

Conway, J. R., Lex, A. and Gehlenborg, N. (2017). UpSetR: an R package for the visualization of intersecting sets and their properties. Bioinformatics 33, 2938–2940.

Cramer, A. A. W., Prasad, V., Eftestøl, E., Song, T., Hansson, K. A., Dugdale, H. F., Sadayappan, S., Ochala, J., Gundersen, K. and Millay, D. P. (2020). Nuclear numbers in syncytial muscle fibers promote size but limit the development of larger myonuclear domains. Nat Commun 11, 6287.

Dehghani-Samani, A., Madreseh-Ghahfarokhi, S. and Dehghani-Samani, A. (2019). Mutations of Voltage-Gated Ionic Channels and Risk of Severe Cardiac Arrhythmias. Acta Cardiol Sin 35, 99–110.

deLapeyrière, O., Ollendorff, V., Planche, J., Ott, M. O., Pizette, S., Coulier, F. and Birnbaum, D. (1993). Expression of the Fgf6 gene is restricted to developing skeletal muscle in the mouse embryo. Development 118, 601–611.

Dos Santos, M., Backer, S., Auradé, F., Wong, M. M., Wurmser, M., Pierre, R., Langa, F., Do Cruzeiro, M., Schmitt, A., Concordet, J. P., et al. (2022). A fast Myosin super enhancer dictates muscle fiber phenotype through competitive interactions with Myosin genes. Nat Commun 13, 1039.

Dos Santos, M., Backer, S., Saintpierre, B., Izac, B., Andrieu, M., Letourneur, F., Relaix, F., Sotiropoulos, A. and Maire, P. (2020). Single-nucleus RNA-seq and FISH identify coordinated transcriptional activity in mammalian myofibers. Nat Commun 11, 5102.

Edom-Vovard, F. and Duprez, D. (2004). Signals regulating tendon formation during chick embryonic development. Dev Dyn 229, 449–457.

Edom-Vovard, F., Schuler, B., Bonnin, M. A., Teillet, M. A. and Duprez, D. (2002). Fgf4 positively regulates scleraxis and tenascin expression in chick limb tendons. Dev Biol 247, 351–366.

Esteves de Lima, J., Blavet, C., Bonnin, M. A., Hirsinger, E., Comai, G., Yvernogeau, L., Delfini, M. C., Bellenger, L., Mella, S., Nassari, S., et al. (2021). Unexpected contribution of fibroblasts to muscle lineage as a mechanism for limb muscle patterning. Nat Commun 12, 3851.

Esteves de Lima, J., Blavet, C., Bonnin, M. A., Hirsinger, E., Havis, E., Relaix, F. and Duprez, D. (2022). TMEM8C-mediated fusion is regionalized and regulated by NOTCH signalling during foetal myogenesis. Development 149.

Esteves de Lima, J., Bonnin, M. A., Birchmeier, C. and Duprez, D. (2016). Muscle contraction is required to maintain the pool of muscle progenitors via YAP and NOTCH during fetal myogenesis. Elife 5.

Esteves de Lima, J. and Relaix, F. (2021). Master regulators of skeletal muscle lineage development and pluripotent stem cells differentiation. Cell Regen 10, 31.

Fabregat, A., Sidiropoulos, K., Viteri, G., Forner, O., Marin-Garcia, P., Arnau, V., D’Eustachio, P., Stein, L. and Hermjakob, H. (2017). Reactome pathway analysis: a high-performance in-memory approach. BMC Bioinformatics 18, 142.

Fernández-Simón, E., Tejedera-Villafranca, A., Fernández-Garabay, X., Clark, J., Monceau, A., Villalobos, E., Cox, D., Azcón, J. R., Fernández-Costa, J. M. and Diaz-Manera, J. (2026). Investigating the role of EGFR signalling in muscle dystrophies: implications for Duchenne muscular dystrophy. Cell Death Dis 17, 18.

Fornasari, B. E., El Soury, M., De Marchis, S., Perroteau, I., Geuna, S. and Gambarotta, G. (2016). Neuregulin1 alpha activates migration of neuronal progenitors expressing ErbB4. Mol Cell Neurosci 77, 87–94.

Garriock, R. J., Meadows, S. M. and Krieg, P. A. (2005). Developmental expression and comparative genomic analysis of Xenopus cardiac myosin heavy chain genes. Dev Dyn 233, 1287–1293.

Gloriani, M., Cheli, B., D’Ercole, C., Ruggieri, V., Cosentino, M., Serrat Pineda, M., Lozanoska-Ochser, B., Grassi, F., Bouché, M., Madaro, L. et al. (2025). Sarcoglycans are enriched at the neuromuscular junction in a nerve-dependent manner. Cell Death Dis 16, 37.

Gonçalves, T. J. M., Boutillon, F., Lefebvre, S., Goffin, V., Iwatsubo, T., Wakabayashi, T., Oury, F. and Armand, A. S. (2019). Collagen XXV promotes myoblast fusion during myogenic differentiation and muscle formation. Sci Rep 9, 5878.

Ham, A. S., Lin, S., Tse, A., Thürkauf, M., McGowan, T. J., Jörin, L., Oliveri, F. and Rüegg, M. A. (2025). Single-nuclei sequencing of skeletal muscle reveals subsynaptic-specific transcripts involved in neuromuscular junction maintenance. Nat Commun 16, 2220.

Hansson, K. A., Eftestøl, E., Bruusgaard, J. C., Juvkam, I., Cramer, A. W., Malthe-Sørenssen, A., Millay, D. P. and Gundersen, K. (2020). Myonuclear content regulates cell size with similar scaling properties in mice and humans. Nat Commun 11, 6288.

Härönen, H., Zainul, Z., Naumenko, N., Sormunen, R., Miinalainen, I., Shakirzyanova, A., Santoleri, S., Kemppainen, A. V., Giniatullin, R., Pihlajaniemi, T. et al. (2019). Correct expression and localization of collagen XIII are crucial for the normal formation and function of the neuromuscular system. Eur J Neurosci 49, 1491–1511.

Hastings, R. L., Boyce, W., Juros, D., Kim, J., Khan, Z., Sethi, A. and Valdez, G. (2025). Loss of COL20A1 in Schwann Cells Alters Regeneration of the Neuromuscular Junction and Locomotor Activity in Mice. J Mol Neurosci 75, 113.

Hirsinger, E., Blavet, C., Bonnin, M. A., Bellenger, L., Gharsalli, T. and Duprez, D. (2024). Limb connective tissue is organized in a continuum of promiscuous fibroblast identities during development. iScience 27, 110305.

Hutcheson, D. A., Zhao, J., Merrell, A., Haldar, M. and Kardon, G. (2009). Embryonic and fetal limb myogenic cells are derived from developmentally distinct progenitors and have different requirements for beta-catenin. Genes Dev 23, 997–1013.

Jiang, C., Liu, T., Dai, Z., Xiang, L., Zhu, Y., Zhou, X., Huang, X., Shen, Y., Liu, J., Ji, Y. et al. (2026). Mechanical unloading promotes adult cardiomyocyte proliferation through epicardial NRG1-ERBB4 signaling. Nat Cardiovasc Res 5, 705–724.

Karlsen, A., Yeung, C. C., Schjerling, P., Denz, L., Hoegsbjerg, C., Jakobsen, J. R., Krogsgaard, M. R., Koch, M., Schiaffino, S., Kjaer, M. et al. (2023). Distinct myofibre domains of the human myotendinous junction revealed by single-nucleus RNA sequencing. J Cell Sci 136.

Kim, I., Ghosh, A., Bundschuh, N., Hinte, L., Petrosyan, E., von Meyenn, F. and Bar-Nur, O. (2022). Integrative molecular roadmap for direct conversion of fibroblasts into myocytes and myogenic progenitor cells. Sci Adv 8, eabj4928.

Kim, M., Franke, V., Brandt, B., Lowenstein, E. D., Schöwel, V., Spuler, S., Akalin, A. and Birchmeier, C. (2020). Single-nucleus transcriptomics reveals functional compartmentalization in syncytial skeletal muscle cells. Nat Commun 11, 6375.

Koch, M., Schulze, J., Hansen, U., Ashwodt, T., Keene, D. R., Brunken, W. J., Burgeson, R. E., Bruckner, P. and Bruckner-Tuderman, L. (2004). A novel marker of tissue junctions, collagen XXII. J Biol Chem 279, 22514–22521.

Korb, A., Tajbakhsh, S. and Comai, G. E. (2024). Functional specialisation and coordination of myonuclei. Biol Rev Camb Philos Soc 99, 1164–1195.

Kowalski, K., Kołodziejczyk, A., Sikorska, M., Płaczkiewicz, J., Cichosz, P., Kowalewska, M., Stremińska, W., Jańczyk-Ilach, K., Koblowska, M., Fogtman, A. et al. (2017). Stem cells migration during skeletal muscle regeneration -the role of Sdf-1/Cxcr4 and Sdf-1/Cxcr7 axis. Cell Adh Migr 11, 384–398.

Kummer, T. T., Misgeld, T. and Sanes, J. R. (2006). Assembly of the postsynaptic membrane at the neuromuscular junction: paradigm lost. Curr Opin Neurobiol 16, 74–82.

L’Honoré, A., Coulon, V., Marcil, A., Lebel, M., Lafrance-Vanasse, J., Gage, P., Camper, S. and Drouin, J. (2007). Sequential expression and redundancy of Pitx2 and Pitx3 genes during muscle development. Dev Biol 307, 421–433.

Lagha, M., Kormish, J. D., Rocancourt, D., Manceau, M., Epstein, J. A., Zaret, K. S., Relaix, F. and Buckingham, M. E. (2008). Pax3 regulation of FGF signaling affects the progression of embryonic progenitor cells into the myogenic program. Genes Dev 22, 1828–1837.

Lejard, V., Blais, F., Guerquin, M. J., Bonnet, A., Bonnin, M. A., Havis, E., Malbouyres, M., Bidaud, C. B., Maro, G., Gilardi-Hebenstreit, P. et al. (2011). EGR1 and EGR2 involvement in vertebrate tendon differentiation. J Biol Chem 286, 5855–5867.

Li, L., Xiong, W. C. and Mei, L. (2018). Neuromuscular Junction Formation, Aging, and Disorders. Annu Rev Physiol 80, 159–188.

Mackey, A. L. (2025). The Myotendinous Junction-Form and Function. Cold Spring Harb Perspect Biol 17.

Macosko, E. Z., Basu, A., Satija, R., Nemesh, J., Shekhar, K., Goldman, M., Tirosh, I., Bialas, A. R., Kamitaki, N., Martersteck, E. M. et al. (2015). Highly Parallel Genome-wide Expression Profiling of Individual Cells Using Nanoliter Droplets. Cell 161, 1202–1214.

Malbouyres, M., Guiraud, A., Lefrançois, C., Salamito, M., Nauroy, P., Bernard, L., Sohm, F., Allard, B. and Ruggiero, F. (2022). Lack of the myotendinous junction marker col22a1 results in posture and locomotion disabilities in zebrafish. Matrix Biol 109, 1–18.

Marics, I., Padilla, F., Guillemot, J. F., Scaal, M. and Marcelle, C. (2002). FGFR4 signaling is a necessary step in limb muscle differentiation. Development 129, 4559–4569.

Messina, G., Biressi, S., Monteverde, S., Magli, A., Cassano, M., Perani, L., Roncaglia, E., Tagliafico, E., Starnes, L., Campbell, C. E. et al. (2010). Nfix regulates fetal-specific transcription in developing skeletal muscle. Cell 140, 554–566.

Munezane, H., Oizumi, H., Wakabayashi, T., Nishio, S., Hirasawa, T., Sato, T., Harada, A., Yoshida, T., Eguchi, T., Yamanashi, Y. et al. (2019). Roles of Collagen XXV and Its Putative Receptors PTPσ/δ in Intramuscular Motor Innervation and Congenital Cranial Dysinnervation Disorder. Cell Rep 29, 4362–4376.e4366.

Murali, M. and MacDonald, J. A. (2018). Smoothelins and the Control of Muscle Contractility. Adv Pharmacol 81, 39–78.

Murphy, M. and Kardon, G. (2011). Origin of vertebrate limb muscle: the role of progenitor and myoblast populations. Curr Top Dev Biol 96, 1–32.

Nahlé, S., Javed, A., Joumier, L., Kherdjemil, Y., Sitolle, J., Khetchoumian, K., Parekh, Y., Krezel, W., Malleshaiah, M., Le Grand, F. et al. (2025). Differentially expressed fusogens specify myocyte states to drive myogenesis. Development 152.

Nicolas, R., Bonnin, M. A., Blavet, C., de Lima, J. E., Legallais, C. and Duprez, D. (2024). 3D-environment and muscle contraction regulate the heterogeneity of myonuclei. Skelet Muscle 14, 27.

Petrany, M. J., Swoboda, C. O., Sun, C., Chetal, K., Chen, X., Weirauch, M. T., Salomonis, N. and Millay, D. P. (2020). Single-nucleus RNA-seq identifies transcriptional heterogeneity in multinucleated skeletal myofibers. Nat Commun 11, 6374.

Rossi, A. C., Mammucari, C., Argentini, C., Reggiani, C. and Schiaffino, S. (2010). Two novel/ancient myosins in mammalian skeletal muscles: MYH14/7b and MYH15 are expressed in extraocular muscles and muscle spindles. J Physiol 588, 353–364.

Schmidt, K., Glaser, G., Wernig, A., Wegner, M. and Rosorius, O. (2003). Sox8 is a specific marker for muscle satellite cells and inhibits myogenesis. J Biol Chem 278, 29769–29775.

Stuart, T., Butler, A., Hoffman, P., Hafemeister, C., Papalexi, E., Mauck, W. M., 3rd, Hao, Y., Stoeckius, M., Smibert, P. and Satija, R. (2019). Comprehensive Integration of Single-Cell Data. Cell 177, 1888–1902.e1821.

Tanaka, T., Wakabayashi, T., Oizumi, H., Nishio, S., Sato, T., Harada, A., Fujii, D., Matsuo, Y., Hashimoto, T. and Iwatsubo, T. (2014). CLAC-P/collagen type XXV is required for the intramuscular innervation of motoneurons during neuromuscular development. J Neurosci 34, 1370–1379.

Tintignac, L. A., Brenner, H. R. and Rüegg, M. A. (2015). Mechanisms Regulating Neuromuscular Junction Development and Function and Causes of Muscle Wasting. Physiol Rev 95, 809–852.

Toulouse, G., Jarassier, W., Jagot, S., Morin, V., Le Grand, F. and Marcelle, C. (2025). Early lineage segregation of primary myotubes from secondary myotubes and adult muscle stem cells. Nat Commun 16, 7858.

Uxa, S., Castillo-Binder, P., Kohler, R., Stangner, K., Müller, G. A. and Engeland, K. (2021). Ki-67 gene expression. Cell Death Differ 28, 3357–3370.

Wakabayashi, T. (2020). Transmembrane Collagens in Neuromuscular Development and Disorders. Front Mol Neurosci 13, 635375.

Wang, H., Noulet, F., Edom-Vovard, F., Tozer, S., Le Grand, F. and Duprez, D. (2010). Bmp signaling at the tips of skeletal muscles regulates the number of fetal muscle progenitors and satellite cells during development. Dev Cell 18, 643–654.

Wickham, H. (2016). ggplot2: Elegant Graphics for Data Analysis. Cham: Springer International Publishing.

Wolock, S. L., Lopez, R. and Klein, A. M. (2019). Scrublet: Computational Identification of Cell Doublets in Single-Cell Transcriptomic Data. Cell Syst 8, 281–291.e289.

Wu, G. and Haw, R. (2017). Functional Interaction Network Construction and Analysis for Disease Discovery. Methods Mol Biol 1558, 235–253.

Yeh, C. J., Nakka, K., Yablonka-Reuveni, Z. and Lepper, C. (2025). FGFR1 and FGFR4 are essential regulators of muscle stem cell quiescence. Cell Commun Signal 23, 535.

Zaben, M. J. and Gray, W. P. (2013). Neuropeptides and hippocampal neurogenesis. Neuropeptides 47, 431–438.

Zappia, L. and Oshlack, A. (2018). Clustering trees: a visualization for evaluating clusterings at multiple resolutions. GigaScience 7.

Zhou, X., Vachon, C., Cizeron, M., Romatif, O., Bülow, H. E., Jospin, M. and Bessereau, J. L. (2021). The HSPG syndecan is a core organizer of cholinergic synapses. J Cell Biol 220.

Zofkie, W., Southard, S. M., Braun, T. and Lepper, C. (2021). Fibroblast growth factor 6 regulates sizing of the muscle stem cell pool. Stem Cell Reports 16, 2913–2927.

