## SupplementaryFigures for "Regionalisation of muscle nucleus heterogeneity in foetal skeletal muscle"

Kurosawa et al.

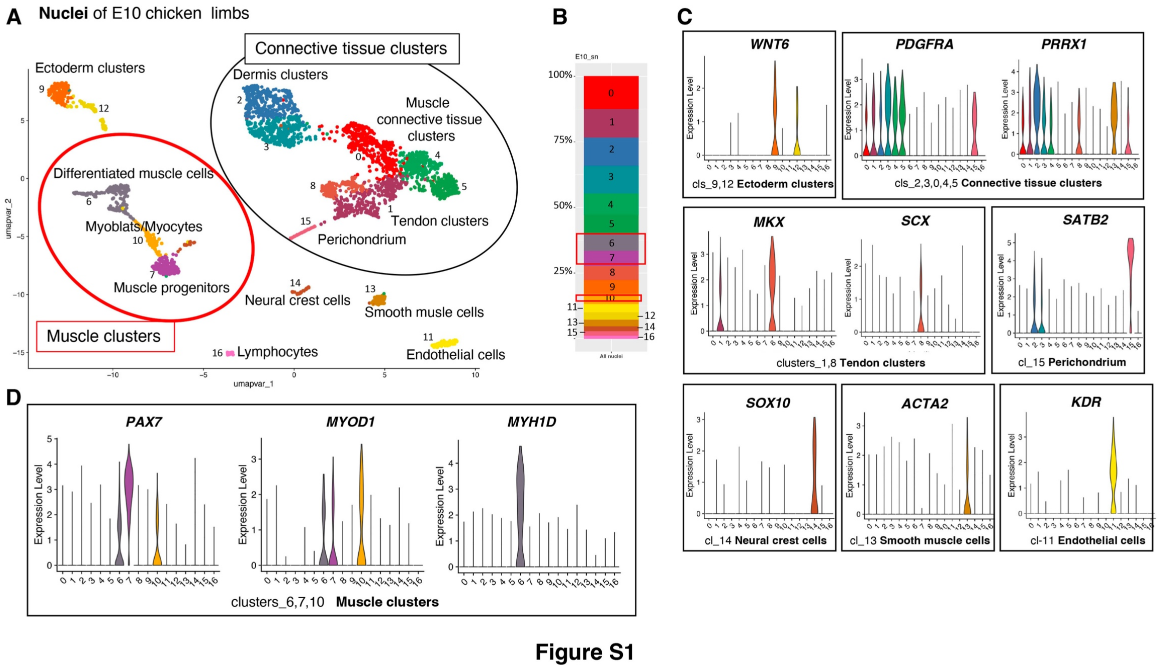


**Figure S1**

**Single-nucleus RNA-sequencing datasets of whole limb nuclei from E10 chicken embryos**

**(A)** UMAP plot showing the distribution of clusters (3174 nuclei) in E10 whole limb snRNAseq dataset. Muscle clusters are circled in red, while connective tissue clusters are circled in black.

**(B)** Stacked histogram showing the percentage of nuclei per cluster in E10 whole limb snRNAseq datasets. Muscle clusters 6,7 and 10 are boxed in red.

**(C)** Violin plots showing the log-normalized expression levels of known markers across clusters in the E10 whole limb snRNAseq dataset, allowing for the identification of the ectoderm clusters (9,12) with *WNT6*, the connective tissue clusters (2,3,0,4,5) with *PDGFRA* and *PRRX1*, the tendon clusters (1,8) with *MKX* and *SCX*, the perichondrium cluster 15 with *SATB2*, the neural crest cluster 14 with *SOX10*, the smooth muscle cluster 13 with *ACTA2* and the endothelial cells cluster 11 with *KDR*.

**(D)** Violin plots showing the log-normalized expression levels of *PAX7*, *MYOD*, *MYH1D* genes across clusters, in E10 whole limb snRNAseq datasets, allowing for the identification of the muscle clusters 6,7,10.

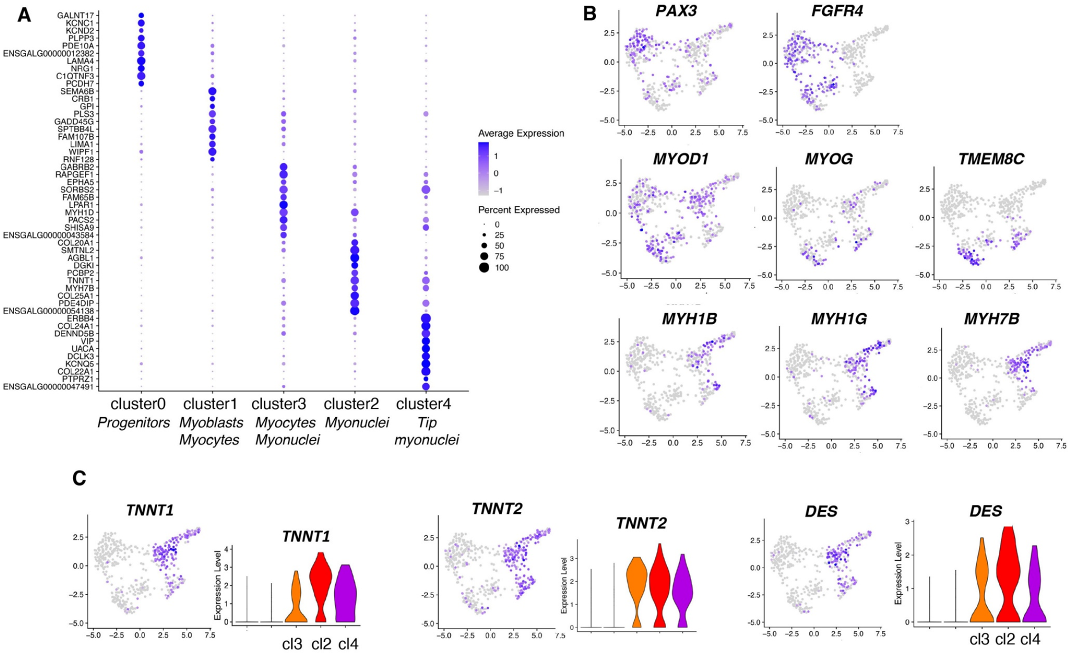


**Figure S2**

#### Figure S2

#### Clustering analysis of the muscle subset of the E10 whole limb snRNAseq dataset

(**A**) Dotplot showing the average expression levels and the fraction of expressing-nuclei for the top10 markers (ordered by fold-change) of each of the five clusters in the E10 muscle snRNAseq dataset, ordered according to myogenic progression.

(**B**) Feature plots showing the log-normalized expression levels across clusters for the following muscle markers, *PAX3*, *FGFR4*, *MYOD1*, *MYOG*, *TMEM8C*, *MYH1B, MYH1G* and *MYH7B* in the E10 muscle snRNAseq datasets Corresponding violin plots are shown in Fig. 2C.

(**C**) Feature and violin plots showing the distribution and log-normalized expression levels for the muscle differentiation genes, *TNNT1, TNNT2* and *DES* across nuclei and clusters.

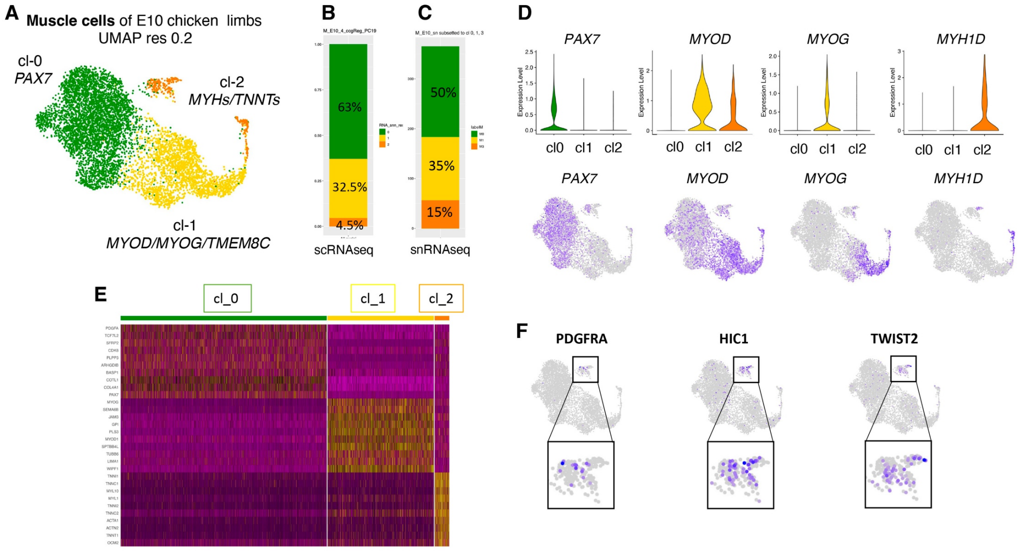


**Figure S3**

### Figure S3

**Myogenic progression illustrated by the analysis of the muscle subset of the E10 whole limb single-cell RNA-sequencing dataset (resolution 0.2)**

(**A**) UMAP plot showing the distribution of the 3 muscle clusters.

**(B)** Stacked histogram showing the percentage of cells per muscle cluster.

(**C**) Stacked histogram showing the percentage of nuclei per cluster for the three first clusters according to myogenic progression (clusters 0, 1 and 3) of the muscle snRNAseq dataset.

(**D**) Violin and feature plots showing the log-normalized expression levels and distribution for the following muscle markers, *PAX7, MYOD1, MYOG* and *MYH1D* across clusters and cells in the E10 muscle scRNAseq dataset at resolution 0.2.

(**E**) Heatmap showing the relative expression of the top 10 markers across all cells of the three clusters (in the muscle scRNAseq dataset at resolution 0.2), ordered according to myogenic progression. Upregulated genes are in yellow, while downregulated genes are in purple.

(**F**) Feature plots with high magnifications showing the log-normalized expression levels for the fibroblast markers *PDGFRA, HIC1* and *TWIST2* across cells pointing to the dual fibroblast/muscle cells.

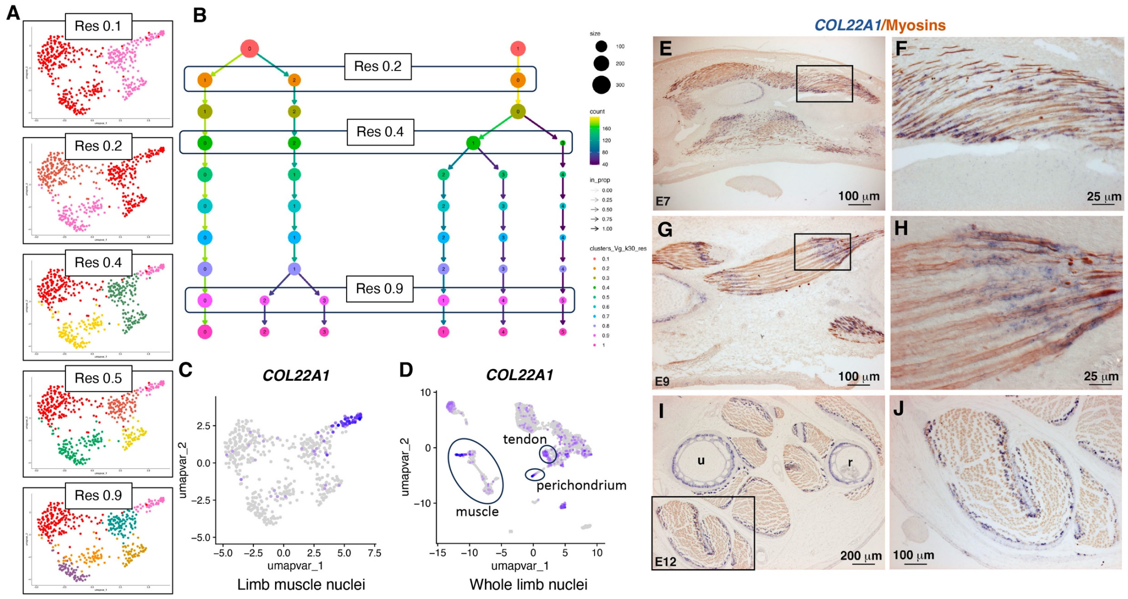


Figure S4

**Figure S4**

***COL22A1* is a reliable marker for tip myonuclei in E10 muscle single-nucleus RNA-sequencing dataset and limb foetal muscles**

(**A**) UMAP plots at different clustering resolutions, 0.1, 0.2, 0.4, 0.5 and 0.9, showing the evolution of the distribution of muscle clusters in the E10 muscle snRNAseq dataset of chicken limb muscles at E10.

(**B**) Clustree tree showing the bifurcations across resolution 0.2, 0.4 and 0.9. The tip myonuclei cluster appears at resolution 0.4 and is observed until resolution 1.

(**C,D**) Feature plots showing the distribution of *COL22A1*+nuclei across clusters in E10 limb muscle (C) and E10 whole limb (D) snRNAseq datasets.

(**E-J**) Colorimetric in situ hybridization to longitudinal (E-H) and transverse (I,J) limb sections for *COL22A1* (blue) followed by an immunostaining with the MF20 antibody to visualize myosins (brown) from E7, E9 and E12 chicken embryos. (F,H,J) are high magnifications of muscles squared in (E,G,I), respectively.

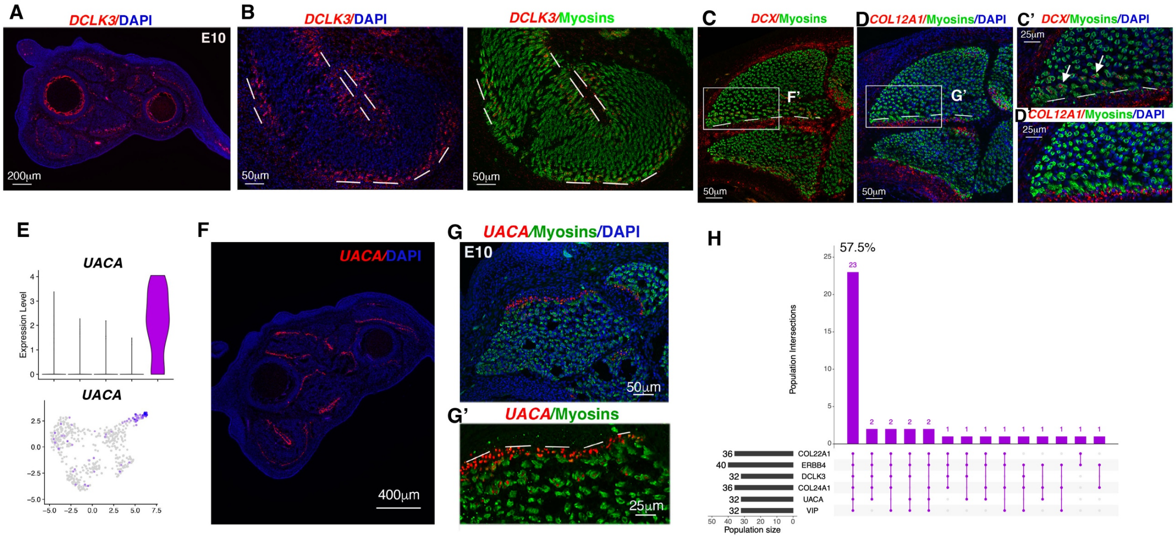


Figure S5

#### Figure S5
#### Molecular markers of tip myonuclei

(**A,B**) Fluorescent in situ hybridization to transverse limb muscle sections of E10 chicken embryos followed with an immunohistochemistry with the MF20 antibody recognising myosins (green) and DAPI staining (blue) with *DCLK3* probe (red). (A) Low magnification of transverse limb section with *DCLK3*/DAPI staining. (B) is a focus of the FCU muscle (ventral and posterior muscle) with *DCLK3/*DAPI staining and *DCLK3*/myosins staining. Dashed lines indicate the muscle/tendon interface.

**(C,D)** Fluorescent in situ hybridization to adjacent transverse limb muscle sections of E10 chicken embryos followed with an immunohistochemistry with the MF20 antibody recognising myosins (green) and DAPI staining (blue) with *DCX* (C) and *COL12A1* (D) probes (red). (C',D') are high magnifications of part of the dorsal muscle squared in upper panels. Dashed lines indicate the muscle/tendon interface. Arrows point to *DCX* expression in tip myonuclei, while *COL12A1* labels tendons.

(**E**) Violin and feature plots showing the log-normalized expression levels and the distribution of the *UACA* gene across clusters of E10 muscle snRNAseq dataset.

(**F,G,G'**) Fluorescent in situ hybridization to transverse limb muscle sections of E10 chicken embryos followed with an immunohistochemistry with the MF20 antibody recognising myosins (green) and DAPI staining (blue) with *UACA* probe (red). (F) Low magnification of transverse limb section with *UACA*/DAPI staining showing the regionalised *UACA* expression in limb muscles. (G) is a focus of a dorsal and posterior muscle showing *UACA* expression in tip myonuclei. Dashed lines indicate the muscle/tendon interface. (G') is a high magnification of the muscle shown in (G) focusing on muscle tip.

(**H**) UpSet plot showing nuclei numbers per gene combination between the top 6 genes of cluster 4 filtered marker list (*COL22A1, ERRB4, DCLK3 COL24A1, UACA, VIP*). The size of each population expressing a gene of interest is shown at the bottom left of the plots.

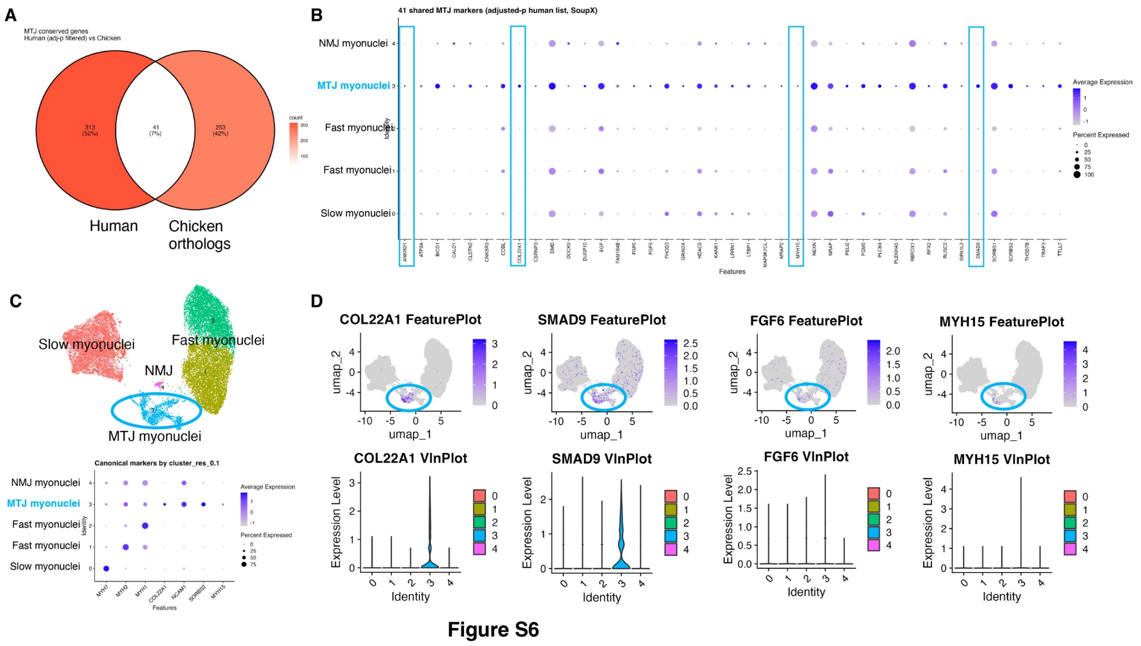


**Figure S6**

**Comparison of the molecular signatures between chicken tip foetal myonuclei with human adult MTJ myonuclei**

(**A**) Venn diagram of MTJ myonuclei genes from human adult muscle snRNAseq datasets (Karlsen et al., 2023) and chicken E10 muscle snRNAseq dataset.

(**B**) Dotplot showing the average expression levels and the fraction of expressing-nuclei for 41 shared genes between the human adult muscle snRNAseq datasets (Karlsen et al., 2023) and the chicken E10 muscle snRNAseq dataset.

(**C**) UMAP plot showing the myonuclei distribution from the three human samples (Karlsen et al., 2023) and dotplot showing the average expression levels and the fraction of expressing-nuclei for the main markers in each of the five clusters of the human adult muscle snRNAseq datasets (Karlsen et al., 2023).

(**D**) Feature and Violin plots showing the distribution and the log-normalized expression levels of four shared genes, *COL22A1, SMAD9, FGF6* and *MYH15* across myonuclei and clusters, in human muscle datasets; genes for which the expression is validated in tip myonuclei of chicken E10 muscles.

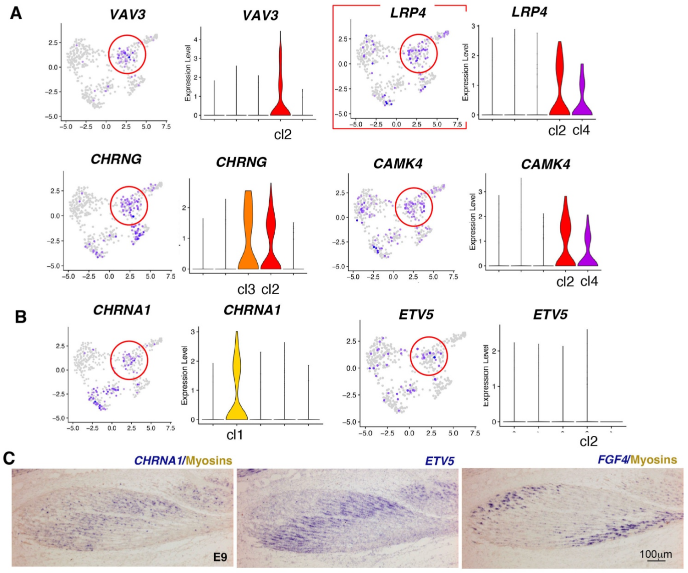


Figure S7

**Figure S7**
**NMJ post-synaptic markers in E10 muscle snRNAseq dataset and in E10 chicken limb muscles**
(**A**) Feature and violin plots showing the distribution and log-normalized expression levels of the *VAV3, LRP4, CHRNG, CAMK4* post-synaptic genes across nuclei and clusters (genes present in non-filtered gene list of cluster 2 and listed by fold-changes high to low)
(**B**) Feature and violin plots showing the distribution and log-normalized expression levels of *CHRNA1* and *ETV5* post-synaptic genes across nuclei and clusters.
(**C**) Colorimetric in situ hybridization to longitudinal muscle sections of E10 chicken limbs for CHRNA1, ETV5 and FGF4 probes (blue), showing the central location of *CHRNA1, ETV5* transcripts and the tip location of *FGF4* transcripts.

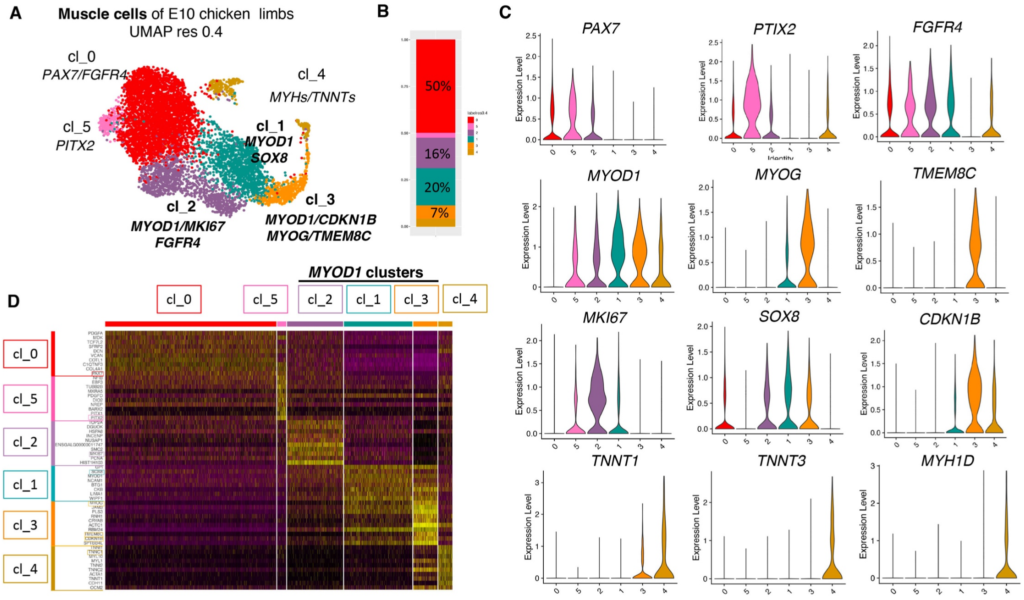


Figure S8

### Figure S8

#### Clustering analysis of the E10 muscle scRNAseq dataset at resolution 0.4

(**A**) UMAP plot showing the distribution of the six muscle clusters at E10, at resolution 0.4, and associated cell composition of each cluster: the two progenitor clusters (0,5), the three MYOD1 clusters (2,1,3) and the myosin cluster 4.

(**B**) Stacked histogram showing the percentage of cells per cluster from the E10 muscle scRNAseq dataset at resolution 0.4.

(**C**) Violin plots showing the log-normalized expression levels for the following muscle markers, *PAX7, PITX2, FGFR4, MYOD1, MYOG, TMEM8C, MKI67, SOX8, CDKN1B, TNNT1, TNNT3* and *MYH1D,* across clusters from the muscle scRNAseq dataset at resolution 0.4.

**(D)** Heatmap showing the relative expression of the top 10 markers across all cells of the six muscle clusters (from the muscle scRNAseq dataset resolution 0.4), ordered according to myogenic progression. Upregulated genes are in yellow, while downregulated genes are in purple.

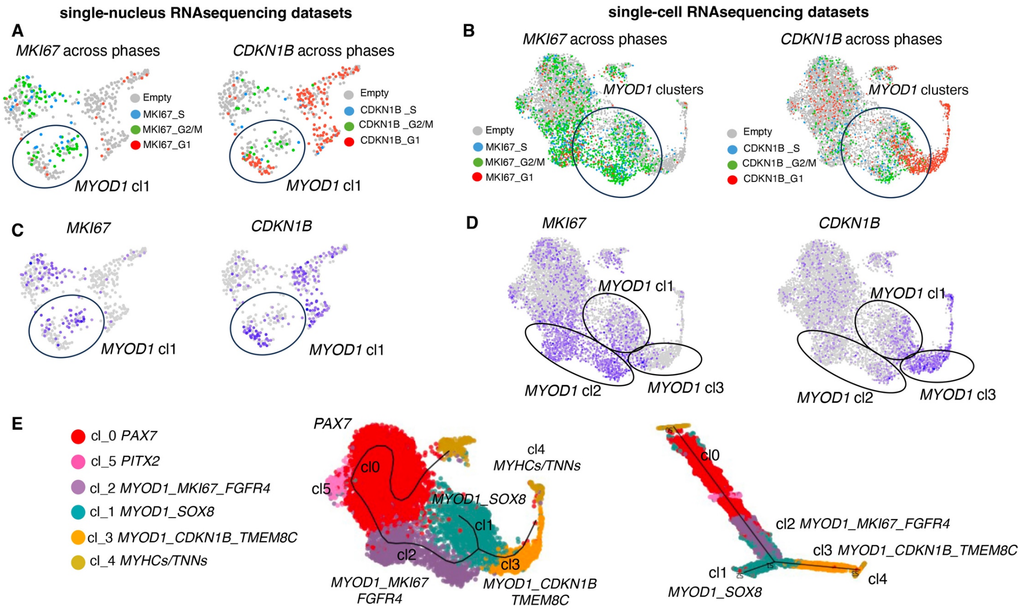


**Figure S9**

**Figure S9**

**Cell cycle markers in the muscle snRNAseq and scRNAseq datasets and STREAM inferred trajectory in the muscle scRNAseq dataset**

(**A,B**) UMAP plot showing the distribution of MKI67+ nuclei/cells and CDKN1B+ nuclei/cells according to cell cycle phases across nuclei/cells in snRNAseq (A) and scRNA seq (B) E10 muscle datasets. *MKI67* is associated with S/G2/M phases, while *CDKN1B* is associated with G1 phase in both datasets.

(**C,D**) Feature plots showing the log-normalized expression levels of *MKI67* and *CDKN1B* across nuclei/cells in E10 muscle snRNAseq (C) and scRNA seq (D) datasets, showing the non-uniform distribution of *MKI67* and *CDKN1B* expression.

(**E**) UMAP and branching tree plot of the inferred trajectory performed on the scRNAseq muscle dataset (res 0.4), showing the distribution of muscle clusters.
